# Ancestral admixture is the main determinant of global biodiversity in fission yeast

**DOI:** 10.1101/415091

**Authors:** Sergio Tusso, Bart P.S. Nieuwenhuis, Fritz J. Sedlazeck, John W. Davey, Daniel C. Jeffares, Jochen B. W. Wolf

**Author notes:** **Corresponding authors:** Sergio Tusso, Jochen B. W. Wolf.

## Abstract

Mutation and recombination are key evolutionary processes governing phenotypic variation and reproductive isolation. We here demonstrate that biodiversity within all globally known strains of *Schizosaccharomyces pombe* arose through admixture between two divergent ancestral lineages. Initial hybridization occurred ~20 sexual outcrossing generations ago consistent with recent, human-induced migration at the onset of intensified transcontinental trade. Species-wide heritable phenotypic variation was explained near-exclusively by strain-specific arrangements of alternating ancestry components with evidence for transgressive segregation. Reproductive compatibility between strains was likewise predicted by the degree of shared ancestry. To assess the genetic determinants of ancestry block distribution across the genome, we characterized the type, frequency and position of structural genomic variation (SV) using nanopore and single-molecule real time sequencing, discovering over 800 SVs. Despite being associated with double-strand break initiation points, SV exerted overall little influence on the introgression landscape or on reproductive compatibility that exist between strains. In contrast, we find strongly increased statistical linkage between ancestral populations that is consistent with negative epistatic selection shaping genomic ancestry combinations during the course of hybridization. This study provides a detailed, experimentally tractable example that genomes of natural populations are mosaics reflecting different evolutionary histories. Exploiting genome-wide heterogeneity in the history of ancestral recombination and lineage-specific mutations sheds new light on the population history of *S. pombe* and highlights the importance of hybridization as a creative force in generating biodiversity.

## Introduction

Mutation is the ultimate source of biodiversity. In sexually reproducing organisms it is assisted by recombination shuffling mutations of independent genomic backgrounds into millions of novel combinations. This widens the phenotypic space upon which selection can act and thereby accelerates evolutionary change (Muller, 1932; Fisher, 1999; McDonald et al., 2016). This effect is enhanced for heterospecific recombination between genomes of divergent populations (Abbott et al., 2013). Novel combinations of independently accumulated mutations can significantly increase the overall genetic and phenotypic variation, even beyond the phenotypic space of parental lineages (transgressive segregation (Lamichhaney et al., 2017; Nolte and Sheets, 2005)). Yet, if mutations of the parental genomes are not compatible to produce viable and fertile offspring, hybridization is a dead end. Phenotypic variation then remains within the confines of genetic variation of each reproductively isolated, parental lineage.

It is increasingly recognised that hybridization is commonplace in nature, and constitutes an important driver of diversification (Abbott et al., 2013; Mallet, 2005). Ancestry components of hybrid genomes can range from clear dominance of alleles from the more abundant species (Dowling et al., 1989; Taylor and Hebert, 1993), over a range of admixture proportions (Lamichhaney et al., 2017; Runemark et al., 2018) to the transfer of single adaptive loci (The Heliconius Genome Consortium et al., 2012). The final genomic composition is determined by a complex interplay of demographic processes, heterogeneity in recombination (e.g. induced by genomic rearrangements) (Wellenreuther and Bernatchez, 2018) and selection (Sankararaman et al., 2014; Schumer et al., 2016). Progress in sequencing technology, now allows characterisation of patterns of admixture and the illumination of underlying processes (Payseur and Rieseberg, 2016). Yet, research has largely focused on animals (Turner and Harr, 2014; Vijay et al., 2016; Meier et al., 2017; Jay et al., 2018) and plants (Twyford et al., 2015) characterized by large genomes and long generation times. Relatively little attention has been paid to natural populations of sexually reproducing micro-organisms (Leducq et al., 2016; Stukenbrock, 2016; Peter et al., 2018; Steenkamp et al., 2018).

The fission yeast *Schizosaccharomyes pombe* is an archiascomycete haploid unicellular fungus with a facultative sexual mode of reproduction. Despite of its outstanding importance as a model system in cellular biology (Hoffman et al., 2015) and the existence of global sample collections, essentially all research has been limited to a single isogenic strain isolated by Leupold in 1949 (Leupold 972; JB22 in this study). Very little is known about the ecology, origin, and evolutionary history of the species (Jeffares, 2018). Global population structure has been described shallow with no apparent geographic stratification (Jeffares et al., 2015). Genetic diversity (π = 3×10^−3^ substitutions/site) appears to be strongly influenced by genome-wide purifying selection with the possible exception of region-specific balancing selection (Fawcett et al., 2014; Jeffares et al., 2015). Despite the overall low genetic diversity, *S. pombe* shows abundant additive genetic variation in a variety of phenotypic traits including growth, stress responses, cell morphology, and cellular biochemistry (Jeffares et al., 2015). The apparent worldwide lack of genetic structure in this species appears inconsistent with the large phenotypic variation between strains and with evidence for post-zygotic reproductive isolation between inter-strain crosses, ranging from 1% to 90 *%* of spore viability (Kondrat’eva and Naumov, 2001; Teresa Avelar et al., 2013; Zanders et al., 2014; Jeffares et al., 2015; Naumov et al., 2015; Marsellach, 2017).

In this study, we integrate whole-genome sequencing data from three different technologies - sequencing-by-synthesis (Illumina technology data accessed from (Jeffares et al., 2015)), single-molecule real-time sequencing (Pacific BioSciences technology, this study) and nanopore sequencing (Oxford Nanopore technology, this study) - sourced from a mostly human-associated, global sample collection to elucidate the evolutionary history of the *S. pombe* complex. Using population genetic analyses based on single nucleotide polymorphism (SNP) we show that global genetic variation and heritable phenotype variation of *S. pombe* results from recent hybridization of two ancient lineages. 25 *de novo* assemblies from 17 divergent strains further allowed us to quantify segregating structural variation including insertions, deletions, inversion and translocations. In light of these findings, we retrace the global population history of the species, and discuss the relative importance of genome-wide ancestry and structural mutations in explaining phenotypic variation and reproductive isolation.

## Results

### Global genetic variation in *S. pombe* is characterized by ancient admixture

Genetic variation of the global *S. pombe* collection comprises 172,935 SNPs segregating in 161 strains. Considering SNPs independently, individuals can be sub-structured into 57 clades that differ by more than 1900 variants, but are near-clonal within clades (Jeffares et al., 2015). To examine population ancestry further, we divided the genome into 1925 overlapping windows containing 200 SNP each and one representative from each clade (57 samples in total). Principle component analysis conducted on each orthologous window showed a highly consistent pattern along the genome (**Figure 1a**, **Supplementary Figure 1**): i) the major axis of variation (PC1) split all samples into two clear discrete groups explaining 60% ±13% of genetic variance (**Figure 1b**). ii) All samples fell into either extreme of the normalized distribution of PC1 scores (*PC*1 ∈ [0; 0.3]x ∪ [0.7; 1]) (**Supplementary Figures 2 & 3**) with the only exception of strains with inferred changes in ploidy level (**Methods**, **Supplementary Figure 4**). iii) PC2 explained 13% ± 6% of variation and consistently attributed higher variation to one of the two groups. This strong signal of genomic windows separating into two discrete groups suggested that the genomic diversity in this collection was derived from two distinct populations. However, iv) group membership of strains changes among windows moving along the genome, reflecting recombination between these two well defined groups. This last point highlights the importance of considering haplotype structure and explains the lack of observed population structure when disregarding non-independence of SNPs (Jeffares et al., 2015).

**Figure 1.**
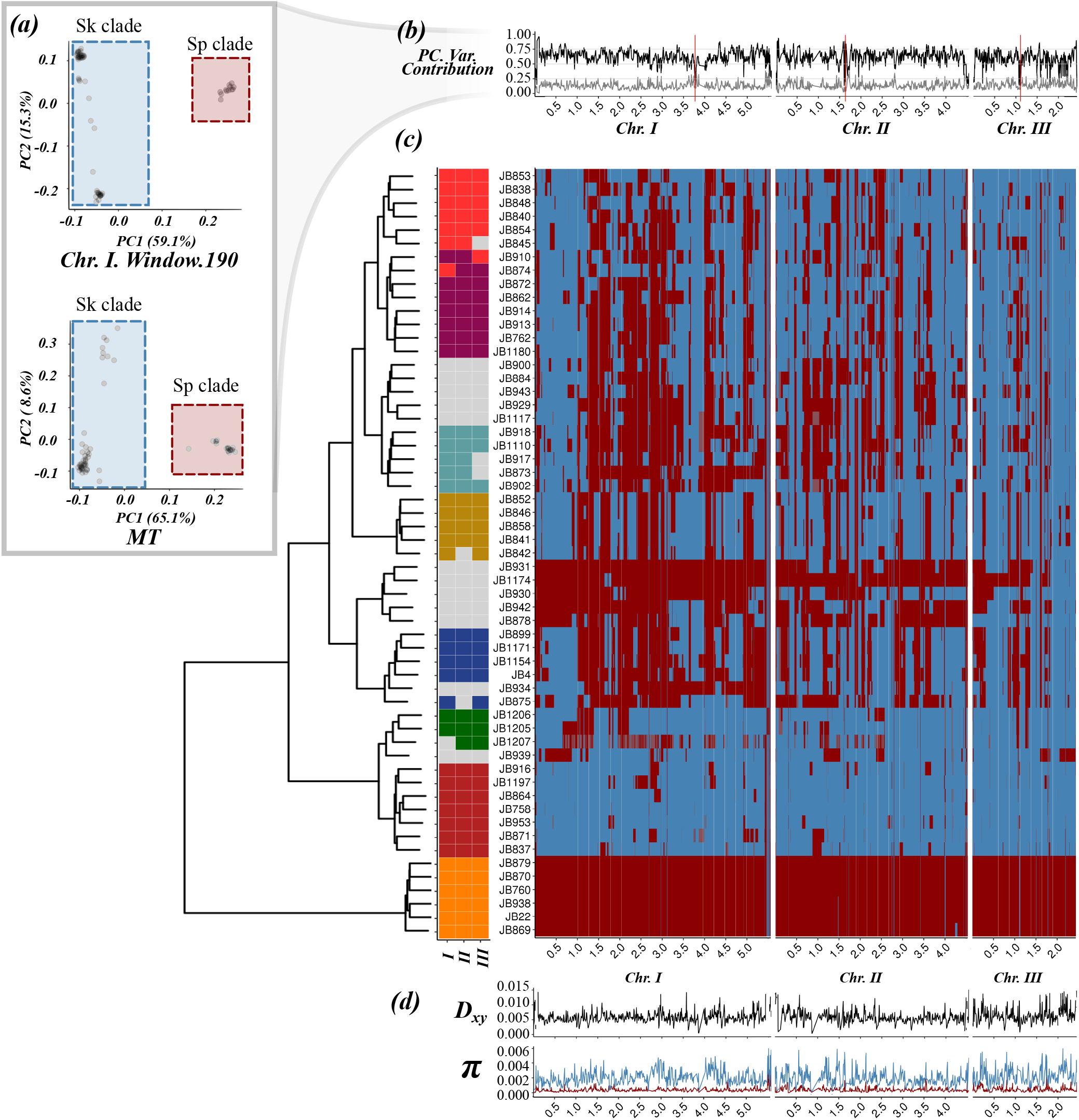
Distribution of *Sp* (red) and *Sk* (blue) ancestry blocks along the *S. pombe genome*. (a) Example of principal component analysis (PCA) of a representative genomic window in chromosome I (top) and the whole mitochondrial DNA (bottom). Samples fall into two major clades, *Sp* (red square) and *Sk* (blue square). The proportion of variance explained by PC1 and PC2 is indicated on the axis labels. Additional examples are found in **Supplementary Figure 1** (b) Proportion of variance explained by PC1 (black line) and PC2 (grey line) for each genomic window along the genome. Centromeres are indicated with red bars. Note the drop in proximity to centromeres and telomeres where genotype quality is significantly reduced. (c) Heatmap for one representative of 57 near-clonal groups indicating ancestry along the genome (right panel). Samples are organised according to a hierarchical clustering, grouping samples based on ancestral block distribution (left dendrogram). Colours on the tips of the cladogram represent cluster membership by chromosome (see **Supplementary Figure 7**). Samples changing clustering group between chromosomes are shown in grey. (d) Estimate of D_xy_ between ancestral groups and genetic diversity (π) within the *Sp* (red) and *Sk* clade (blue) along the genome.

The strong signal from the PCA that systematically differentiates between groups along the genome were likewise reflected in population genetic summary statistics including Watterson’s theta (θ), pairwise nucleotide diversity (π), and Tajima’s D (**Figure 1d and 2**). Significant differences in these statistics (Kendall’s τ p-value ≤ 2.2×10^−16^), were also present in mitochondrial genetic variation (**Figure 1a**), allowed polarising the two groups across windows into a “low-diversity” group (red) and a “high-diversity group (blue) (**Figure 1a**, **Supplementary Figure 5**). Genetic divergence between groups (D_xy_) was 15 and 3 times higher than mean genetic diversity (π) within each group, respectively, and thus supports a period of independent evolution. Painting genomic windows by group membership revealed blocks of ancestry distributed in sample specific patterns along the genome (**Figure 1c**, **Supplementary Figure 6**). The sample corresponding to the reference genome isolated originally from Europe (Leupold’s 972; JB22) consisted almost exclusively of ‘red’ ancestry (>96% red), whereas other samples were characterized near-exclusively by “blue” ancestry (>96% blue). The sample considered to be a different species from Asia, *S. kambucha* (JB1180 (Singh and Klar, 2002)) had a large proportion of “blue” windows (>70% blue). Hereafter, we refer to the “red” and “blue” clade as *Sp* and *Sk*, for *S. pombe* and *S. kambucha* respectively. Grouping samples by the pattern of genomic ancestry across the genome revealed 8 discrete clusters (**Figure 1c**). Consistent with independent and/or recent segregation of ancestral groups, cluster membership for several samples differed between chromosomes (**Figure 1c**) and genome components (**Supplementary Figures 7 & 8**). This is also reflected by low support in the relationship between the 8 discrete clusters.

**Figure 2.**
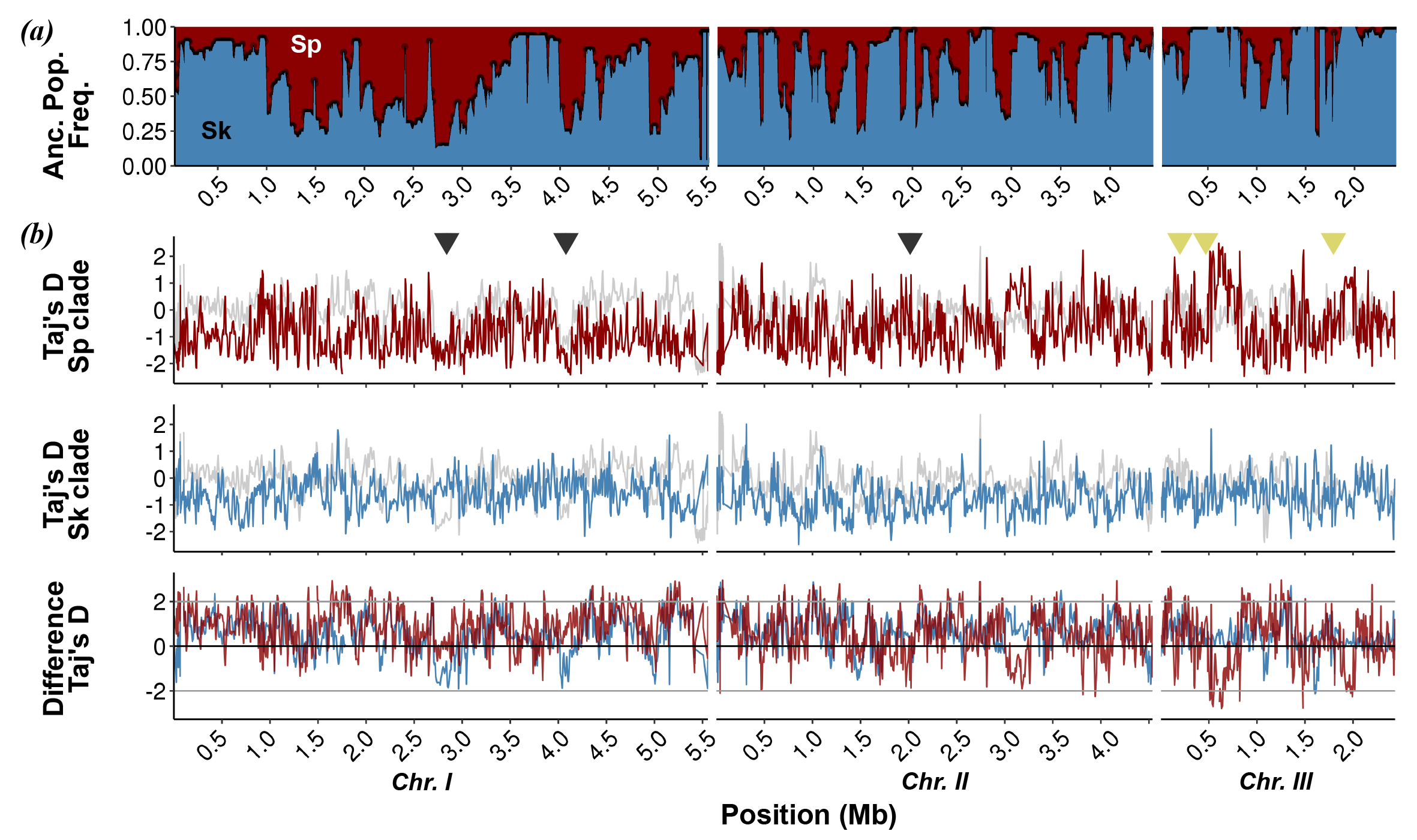
Population genetic summary statistics. (a) Proportion of *Sp* (red) and *Sk* (blue) ancestry across all 57 samples along the genome. (b) Tajima’s D differentiated by *Sp* (red) and *Sk* (blue) ancestry and pooled across all samples irrespective of ancestry (grey line). Genomic regions previously identified under purifying selection (Fawcett et al., 2014) are shown with black triangles. Reported active meiotic drives (Zanders et al., 2014; Hu et al., 2017; Nuckolls et al., 2017) are indicated by yellow triangles. The third panel shows the difference between ancestry specific Tajima’s D and the estimate from the pooled samples.

The distribution of ancestry components was highly heterogeneous across the chromosome (**Figure 2a**, **Supplementary Figure 6**). Most strains showed an excess of *Sp* ancestry in parts of chromosome I, whereas several regions of chromosome III had an excess of *Sk* ancestry. Failing to incorporate this genome-wide variation of admixture proportions can mimic signatures of selection. For example, equal ancestry contributions for a certain genomic region will yield high positive values of both Tajima’s D (**Supplementary Figure 9**) and π and may be mistaken as evidence for balancing selection. Strong skew in ancestry proportions reduces both statistics to values of the prevailing ancestry and may appear as evidence for selective sweeps (**Figure 2b**). Taking ancestry into account, however, there was no clear signature of selection in either *Sp* or *Sk* genetic variation that could account for heterogeneity in the genetic composition of hybrids (**Supplementary Figure 9**). Signatures of selection identified previously (cf. Fawcett et al., 2014) are likely artefacts due to skewed ancestry proportions rather than events of positive or balancing selection in the ancestral populations.

Overall, our results provide strong evidence for the presence of at least two divergent ancestral populations: one genetically diverse group (*Sk* clade) and a less diverse group (*Sp* clade). We found a large range of ancestral admixture proportions between these two clades broadly clustering samples into 8 weakly supported groups. These resemble clusters of strains previously identified by *Structure* and *fineStructure* without explicit modelling of ancestral admixture (Jeffares et al., 2015). Neglecting ancestry, Jeffares et al. (2015) argued that the shallow population structure likely results from extensive gene flow between clusters. Yet, considering the genome-wide distribution of *Sk* and *Sp* ancestry, and lack of geographical structure, suggest that the 8 clusters are derived from one or a few centres of ancient admixture (hybridization) without the need of subsequent or recent gene flow between them.

### Age of ancestral lineages and timing of hybridization

To shed further light on the population history, we estimated the age of the parental lineages and the timing of initial hybridization. Calibrating mitochondrial divergence by known collection dates over the last 100 years, Jeffares et al. (2015) estimated that the time to the most recent common ancestor for all samples was around 2,300 years ago. Current overrepresentation of near-pure *Sp* and *Sk* in Europe or Africa / Asia, respectively, is consistent with an independent history of the parental lineages on different continents for the most part of the last millennia (**Supplementary Figure 8**). Yet, the variety of admixed genomes bears testimony to the fact that isolation has been disrupted by heterospecific gene flow. Using a theoretical model assuming secondary contact with subsequent hybridization (Janzen et al., 2018) we estimated that hybridization occurred within the last 20 sexual outcrossing generations (**Figure 3**, **Supplementary Figures 10 & 11**). Considering intermittent generations of asexual reproduction, high rates of haploid selfing and dormancy of spores (Farlow et al., 2015; Jeffares, 2018) it is difficult to obtain a reliable estimate of time in years. This recent estimate of hybridization is consistent with hybridization induced by the onset of regular trans-continental human trade between Europe with Africa and Asia (~14th century) and with the Americas (~16th century), with fission yeast as a human commensal (Jeffares, 2018). This fits with the observation that all current samples from the Americas were hybrids, while samples with the purest ancestry stem from Europe, Africa and Asia. Moreover, negative genome-wide Tajima’s D estimates for both ancestral clades (mean ± SD for Sp: −0.8 ± 0.9 and Sk: −0.7 ± 0.6) signal a period of recent expansion.

**Figure 3.**
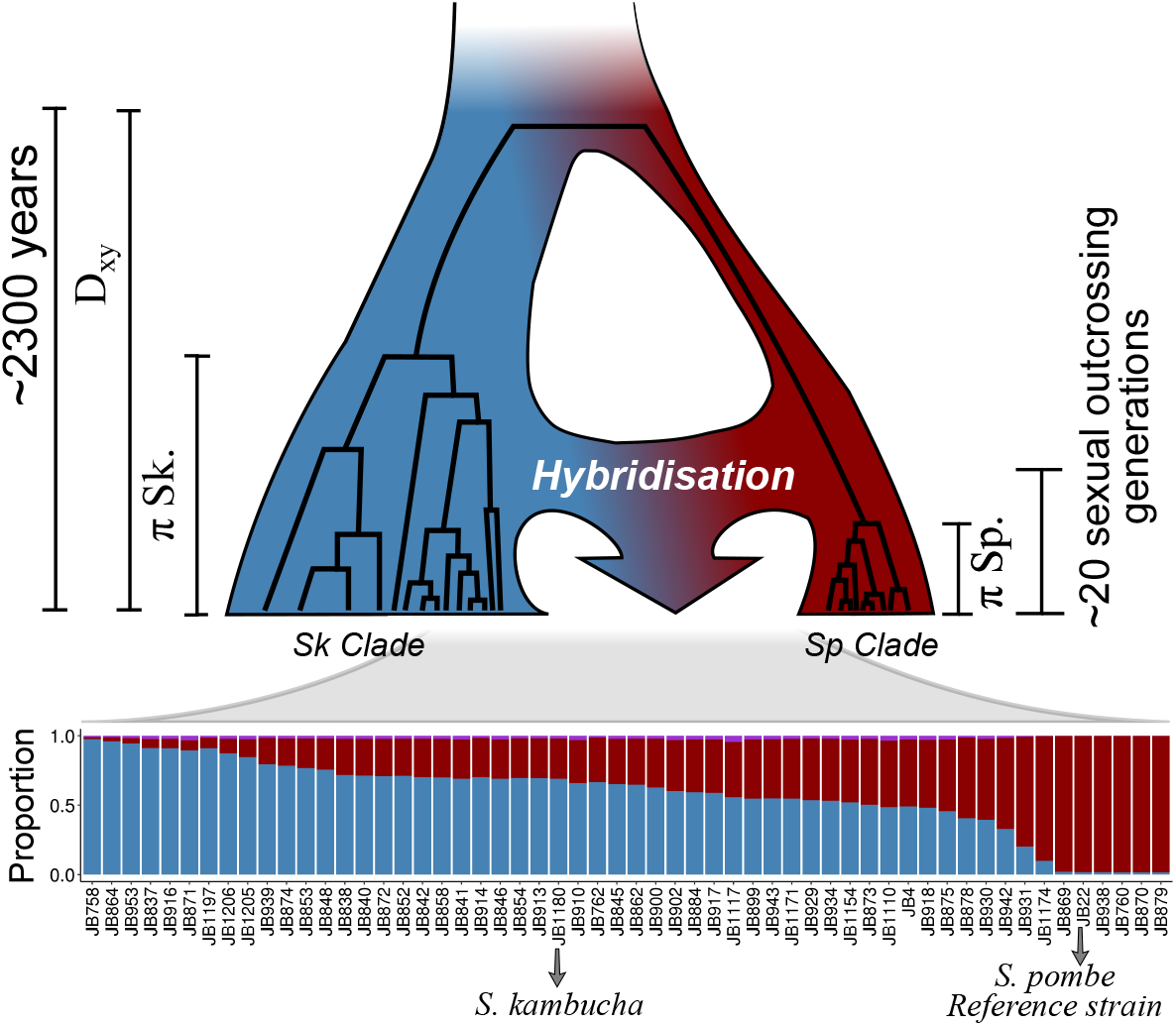
Inferred evolutionary history of contemporary *S. pombe* strains. An ancestral population diverged into two major clades, *Sp* (red) and *Sk* (blue) since approximately 2300 years ago (Jeffares et al., 2015). Recurrent hybridization upon secondary contact initiated around 20 sexual outcrossing generations ago resulted in admixed genomes with a range of admixture proportions (bottom) prevailing today.

### Heritable phenotypic variation and reproductive isolation are governed by ancestry components

Hybridization can lead to rapid evolution due to selection acting on the genetic and phenotypic variation emerging after admixture (Muller, 1932; Fisher, 1999; McDonald et al., 2016). We assessed the consequences of hybridization on phenotypic variation making use of a large data set including 228 quantitative traits collected from the strains under consideration here (Jeffares et al., 2015). Contrary to genetic clustering of hybrid genomes (**Figure 1c**), samples with similar ancestry proportions did not group in phenotypic space described by the first two PC-dimensions capturing 31% of the total variance across traits (**Figure 4a**). Moreover, phenotypic variation of hybrids exceeded variation of pure strains (>0.9 ancestry for *Sp* or Sk). This was supported by trait specific analyses. We divided samples into three discrete groups: pure *Sp*, pure *Sk* and hybrids with a large range of *Sp* admixture proportions (0.1-0.9). 63 traits showed significant difference among groups (**Figure 4b**, **Supplementary Figure 12**). In the vast majority of cases (50 traits), hybrid phenotypes were indistinguishable from one of the parents, but differed from the other, suggesting dominance of one ancestral background, consistent with some ecological separation of the backgrounds. In seven traits, hybrid phenotypes were intermediate differing from both parents, indicative of an additive contribution of both ancestral backgrounds. For six traits, hybrids exceeded phenotypic values of both parents providing evidence for transgressive segregation. In all cases, the number of significantly differentiated traits was found to be higher than under the null model (mean number of significant traits after 10000 permutations: dominant Sk 4 +/- 2, dominant Sk 4 +/- 2, transgressive 0 +/- 0.3, intermediate 0 +/- 0.1; **Supplementary Figure 13**). Jeffares et al. (2017) showed that for each trait the total proportion of phenotypic variance explained by the additive genetic variance component (used as an estimated of the narrow-sense heritability) ranged from 0 to around 90%. We found that across all 228 traits, considering *Sp* and *Sk* ancestry components across the 1,925 genomic windows explained an equivalent amount of phenotypic variance as all 172,935 SNPs segregating across all samples, being both highly correlated (**Figure 4c, 4d**; *r* = 0.82, *p*-value ≤ 2.2 · 10^−16^). Combinations of ancestral genetic variation appear to be the main determinants of heritable phenotypic variation with only little contribution from single-nucleotide mutations arising after admixture. In turn, this supports that the formation of hybrids is recent (see above), and few (adaptive) mutations have occurred after it.

**Figure 4.**
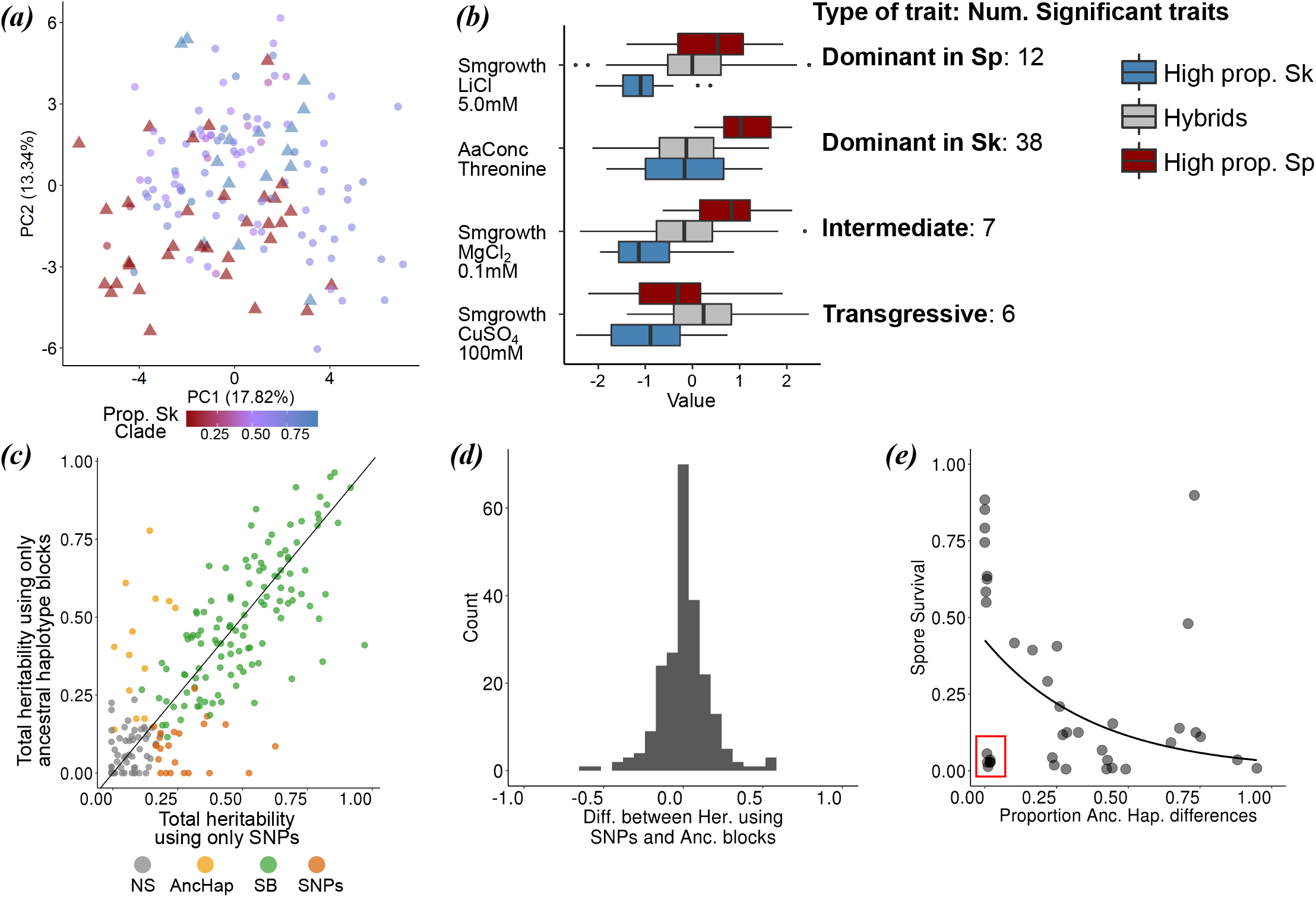
Ancestry explains variation in phenotype and reproductive isolation. (a) PCA of normalized phenotypic variation across 228 traits. The proportion of variance explained by PC1 and PC2 is indicated on the axis labels. Admixed samples (dots) are coloured coded by ancestry proportion (cf. **Figure 3**) ranging from pure *Sp* (red triangle) to pure *Sk* (blue triangle) ancestry. (b) Phenotypic distribution of example traits separated by the degree of admixture: admixed samples are shown in grey, pure ancestral *Sp* and *Sk* samples are shown in red and blue respectively. The number of traits corresponding to a dominant, additive and transgressive genetic architecture is indicated on the right hand side (c) Comparison of heritability estimates of all 228 traits based on 172,935 SNPs (abscissa) and on 1925 genomic windows polarized by ancestry (ordinate). Colours indicate statistical significance. *NS*: heritability values not significantly different from zero, *AncHap:* significant only using ancestral blocks, *SNPs:* significant only using SNPs, *SB:* significant in both analyses. Diagonal (slope=1) added as reference. (d) Histogram of the difference between heritability estimates using SNPs and ancestry components for all 228 traits. (e) Correlation between the difference in ancestry proportions between two strains (cf. **Figure 2**) and spore viability of the cross. Red box shows samples with low spore viability but high genetic similarity.

Ancestry also explained most of the variation in postzygotic reproductive isolation between strains. Previous work revealed a negative correlation between spore viability and genome-wide SNP divergence between strains (Jeffares et al., 2015). The degree of similarity in genome-wide ancestry had the same effect: the more dissimilar two strains were in their ancestry, the lower the viability of the resulting spores (**Figure 4e**; Kendall correlation coefficients, τ = −0.30, T= 259, p-value = 6.66 · 10^−3^). This finding is consistent with reproductive isolation being governed by many, genome-wide incompatibilities between the *Sp* and *Sk* clade. Yet, in a number of cases spore survival was strongly reduced in strain combinations with near-identical ancestry. In these cases, reproductive isolation may be caused by few large effect mutations, including structural genomic changes that arose after hybridization.

### Structural mutations do not determine the genome-wide distribution of ancestry blocks

Structural genomic changes (structural variants or SVs hereafter) are candidates for large-effect mutations governing phenotypic variation (Küpper et al., 2016; Jeffares et al., 2017), reproductive isolation (Hoffmann and Rieseberg, 2008; Teresa Avelar et al., 2013) and heterospecific recombination (Ortiz-Barrientos et al., 2016). They may thus importantly contribute to shaping heterogeneity in the distribution of ancestry blocks observed along the genome (Jay et al., 2018; Poelstra et al., 2014) (**Figure 2b**). However, inference of SVs in natural strains of fission yeast has been primary based on short-read sequencing (Jeffares et al., 2017). SV calls from short-read sequencing data are known to differ strongly by bioinformatic pipeline, are prone to false positive inference and are limited in their ability to infer long-range SV, in particular in repetitive regions of the genome (Jeffares et al., 2017).

To obtain a reliable and comprehensive account of SV segregating across strains and test for a possible association of SVs with the skewed ancestry in the genome, we generated chromosome-level *de novo* genome assemblies for 17 of the most divergent samples using single-molecule real time sequencing (mean sequence coverage 105x; **Supplementary Table 7**). For the purpose of methodological comparison, we also generated *de novo* assemblies for a subset of 8 strains (including the reference Leupold’s 972) based on nanopore sequencing (mean sequence coverage: 140x). SVs were called using a mixed approach combining alignment of *de novo* genomes and mapping of individual reads to the reference genome (Wood et al., 2002). Both approaches and technologies yielded highly comparable results (**Methods**, **Supplementary Figure 14-17** and **Supplementary Table 8**).

After quality filtering, we retained a total of 832 variant calls including 563 insertions or deletions (indels), 118 inversions, 110 translocations and 41 duplications. The 17 strains we examined with long reads could be classified into six main karyotype arrangements (**Figure 5a**). The previously reported list of SVs of the same strains using short reads consisted of only 52 SVs (Jeffares et al., 2017) of which only 8 were found to overlap with the 832 calls from long-read data. The vast majority of SVs were smaller than 10 kb (**Figure 5b**). The size distribution was dominated by elements of 6 kb and 0.5 kb in length corresponding to known transposable elements (TEs) and their flanking long terminal regions (LTRs), respectively (Kelly and Levin, 2005). Only a small number of SVs corresponded to large-scale rearrangements (50 kb - 2.2 Mb) including translocations between chromosomes (**Figure 5a**). A subset of these have been characterized previously as large-effect modifiers of recombination promoting reproductive isolation (Brown et al., 2011; Teresa Avelar et al., 2013; Jeffares et al., 2017).

**Figure 5.**
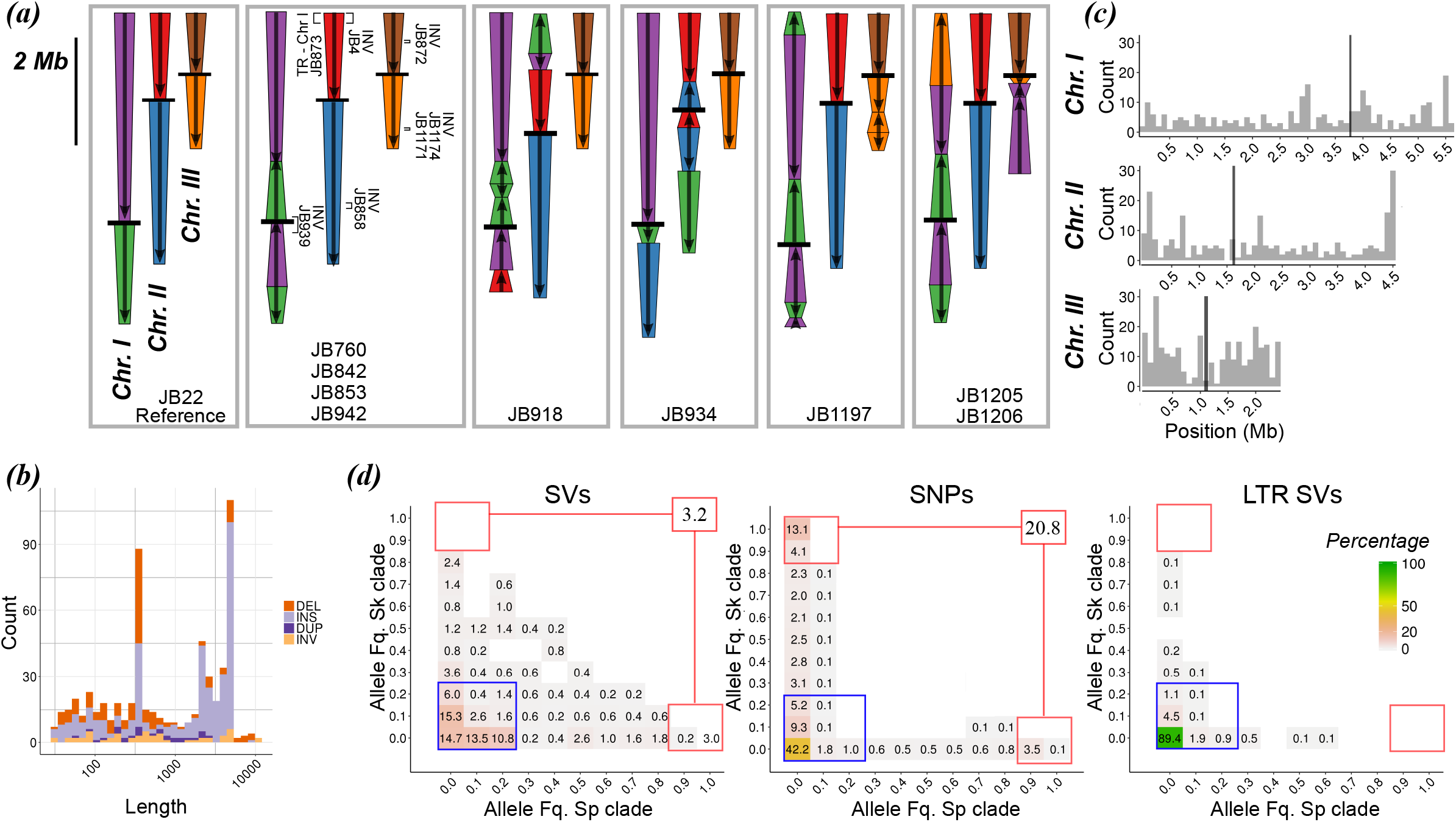
Characterization of structural variation based on long-read, real-time sequencing. (a) Schematic representation of the three chromosomes in different strains displaying SVs larger than 10kb relative to reference genome JB22 (left panel). Chromosome arms are differentiated by colour; orientation is indicated with arrows relative to the reference; black bars represent centromeres. In the second panel, additional SVs, their type and ID of the corresponding strain are illustrated in brackets. (b) Size distribution of SVs below 10 kb. Colours indicate the type of SV. (c) Distribution of SV density along the genome. Black bars represent centromeres. (d) Two-dimensional, folded site frequency spectrum between inferred ancestral populations for all SVs, SNPs and LTR INDELs. Numbers and colours show the percentage of the total number of variants in each category. Variants with low frequency in both populations are shown in the blue box. Variants highly differentiated between populations are show in red boxes with total in the upper right box. Fills with percentage lower than 0.01 are empty. *Abbreviations:* DEL: deletion; DUP: duplication; INS: insertion; INV: inversion

Contrary to previous SV classification based on short reads (Jeffares et al., 2017), SV density was not consistently increased in repetitive sequences such as centromeric and telomeric regions illustrating the difficulty of short-read data in resolving SV in repetitive regions (**Figure 5c**). Instead, we found that the frequency of SV was significantly elevated in close proximity to developmentally programmed DNA double-strand breaks (DSB) associated with recombination initiation (Fowler et al., 2014). The proportion of SV observed within [0, 0.5) kb and [0.5, 1) kb of DSB was increased by 46% (p-value<1×10^−04^) and 67% (p-value<1×10^−04^) relative to random expectations. On the contrary, regions more distant than 10 kb from DSB were relatively depleted of SV (**Supplementary Figure 18**).

Next, we imputed the ancestry of SV alleles from SNPs surrounding SV break points. We calculated allele frequencies for SV in both ancestral clades and constructed a folded two-dimensional site frequency spectrum (**Figure 5d**). The majority of variants (66 %) segregated at frequencies below 0.3 in both ancestral genetic backgrounds. Very few SVs were differentiated between ancestral populations (3 *%* of variants with frequency higher than 0.9 in one population and below 0.1 in the other). This pattern contrasted with the reference spectrum derived from SNPs where the proportion of low frequency variants was similar at 60 %, but genetic differentiation between populations was substantially higher (21 % of SNP variants with frequency higher than 0.9 in one population and below 0.1 in the other). The difference was most pronounced for large SVs (larger than 10 kb) and TEs, for which we estimated allele frequencies for all 57 strains by means of PCR and short-read data, respectively. For TE’s, 98 % of the total 1048 LTR variants segregated at frequencies below 0.3 in both ancestral populations without a single variant differentiating ancestral populations (**Figure 5d**). Large SVs likewise segregated at low frequencies, being present at most in two strains out of 57. This included the translocation reported for *S. kambucha* between chromosome II and III (Zanders et al., 2014), which we found to be specific for that strain. Only the large inversion on chromosome I segregated at higher frequency being present in five strains out of 57, of which three were of pure *Sp* ancestry including the reference strain (**Supplementary Table 10**). Additionally, SV segregating at high frequency (> 0.7) tended to cluster in genomic regions with steep transitions in ancestry between *Sp* and *Sk* ancestry (p-value > 0.1; **Supplementary Figure 19**).

In summary, long-read sequencing provided a detailed account of species-wide diversity in structural genetic variation including over 800 high-quality variants ranging from small indels to large-scale inter-chromosomal rearrangements. SV calls showed substantial overlap among technologies (Pacific Biosciences, Nanopore) and approaches (de novo assembly vs. mapping), but less than 1 % of this variation was inferred from short-read data. This finding admonishes to caution when interpreting SV calls from short read data that is moreover sensitive to genotyping methods. In contrast to genome-wide SNPs, SVs segregated near-exclusively at low frequencies and were rarely differentiated by ancestral origin. This is consistent with strong diversity-reducing purifying selection relative to SNPs. The fact that SVs, including large-scale rearrangements with known effects on recombination and reproductive isolation (Brown et al., 2011; Teresa Avelar et al., 2013; Zanders et al., 2014), were often unique to single strains precludes a role of SVs in shaping patterns of ancestral heterospecific recombination. Moreover, while being concentrated in proximity to double-strand breaks, possibly due to improper repair upon recombination (Currall et al., 2013), SV were not significantly associated with steep transitions in ancestry blocks. Summarizing the evidence, SV appears to have had little influence in shape genome-wide patterns of ancestral admixture and cannot explain the prevalence of reproductive isolation as a function on ancestral similarity (**Figure 4e**).

### Negative epistasis shapes the distribution of ancestral blocks

Alternatively, heterogeneity in the distribution and frequency of ancestry along the genome may result from negative epistatic interactions of incompatible genetic backgrounds (Schumer et al., 2016). An excess of homospecific combinations of physically distant loci can serve as an indication of epistatic selection against genetic incompatibilities which can be segregating at appreciable frequencies even within species (Corbett-Detig et al., 2013). We tested this hypothesis by measuring ancestry disequilibrium (AD) between all possible pairs of genomic windows within a chromosome. Specifically, we quantified linkage disequilibrium (LD) between windows dominated by alleles from the same ancestral group (> 0.7) *Sp-Sp* or *Sk-Sk* (reflecting positive AD) and contrasted it to the degree of linkage disequilibrium arising between heterospecific allele combinations *Sp-Sk* (negative AD) (**Supplementary Figure 20**). LD differed significantly between these two cases (**Figure 6**). While negative AD decreased rapidly with genetic distance (R^2^ < 0.2 after 66, 19 and 21 kb respectively for each chromosome) positive LD was higher in magnitude and extended over larger distances (R^2^ < 0.2 after 1.02, 0.54, and 0.18 Mb respectively for each chromosome in *Sk-Sk* comparisons and 1.59, 1.12, and 0.32 Mb for *Sp-Sp* comparisons). These results are consistent with a role of epistatic selection during the course of hybridization shaping the ancestry composition of admixed genomes.

**Figure 6.**
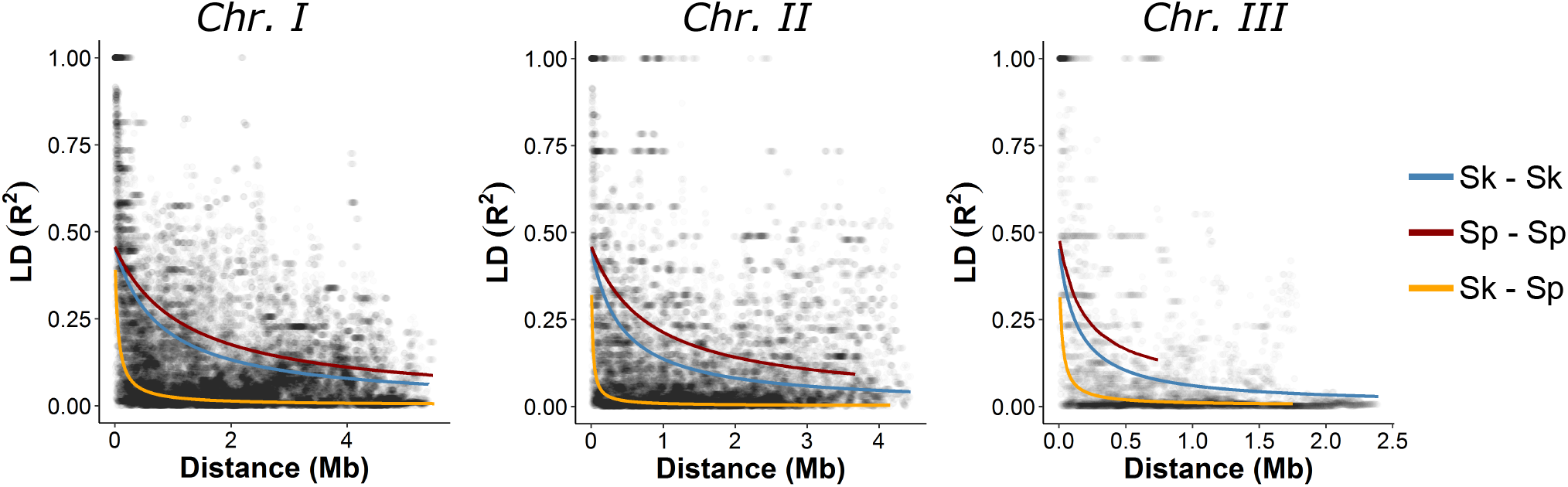
Decay in linkage disequilibrium (LD) with genetic distance. Relationship between LD (R^2^) and physical distance is depicted for each chromosome. Black points represent values for each window pair comparison. Lines show non-lineal regression model based on Hill & Weir (1988) and Remington et al. (2001). LD estimates were divided into three categories representing comparison between windows of shared ancestry (*Sp-Sp* or *Sk-Sk*) reflecting positive ancestry disequilibrium (AD) or of opposite ancestry (*Sk-Sp*) reflecting negative AD.

## Discussion

This study adds to the increasing evidence that hybridization plays an important role as a rapid, “creative” evolutionary force in natural populations (Seehausen, 2004; Mallet, 2007; Soltis and Soltis, 2009; Abbott et al., 2013; Schumer et al., 2014; Abbott et al., 2016; Pennisi, 2016; Nieto Feliner et al., 2017). Recent heterospecific recombination between two ancestral *S. pombe* populations shuffled genetic variation of genomes that diverged since classical antiquity about 2,300 years ago. The timing of hybridization coincided with the onset of intensified trans-continental human trade, suggesting an anthropogenic contribution. Several samples showed similar distribution of ancestral blocks along the genome suggesting comparable evolutionary histories, and allowing the identification of 8 discrete clusters. These clusters, in general showed weak geographical grouping, initially interpreted as evidence for reduced population structure with large recent world-wide gene flow (Jeffares et al., 2015). In contrast, the world-wide distribution of the two ancestral linages suggests rapid and recent global dispersion after hybridization followed by local differentiation. This study thus highlights the importance of taking genomic non-independence into account. Allowing for the fact that genomes are mosaics reflecting different evolutionary histories can fundamentally alter inference on a species’ evolutionary history.

Moreover, conceptualizing genetic variation as a function of ancestry blocks alternating along the genome changes the view on adaptation. Admixture is significantly faster than evolutionary change solely driven by mutation. Accordingly, phenotypic variation was near-exclusively explained by ancestry components with only little contribution from novel mutations. Importantly, admixture not only filled the phenotypic space between parental lineages, but also promoted transgressive segregation in several hybrids. This range of phenotypic outcomes opens the opportunity for hybrids to enter novel ecological niches (Nolte and Sheets, 2005; Pfennig et al., 2016) and track rapid environmental changes (Eroukhmanoff et al., 2013).

Structural mutations have been described as prime candidates for rapid large-effect changes with implications on phenotypic variation, recombination and reproductive isolation (Faria and Navarro, 2010; Ortiz-Barrientos et al., 2016; Wellenreuther and Bernatchez, 2018). This study contributes to this debate providing a detailed account of over 800 high-quality structural variants identified across 17 chromosome level *de nov*o genomes sampled from the most divergent strains within the species. On the whole, SVs had little effect. While large-scale rearrangements in specific strains have been shown to affect fitness (Teresa Avelar et al., 2013; Nieuwenhuis et al., 2018) and promote reproductive isolation between specific strains in *S. pombe* (Brown et al., 2011; Teresa Avelar et al., 2013), reproductive isolation was overall best predicted by the degree of shared ancestry with little contribution from SVs. SVs segregated at low frequencies in both ancestral populations and, contrary to what has been suggested for specific genomic regions in other systems (Jay et al., 2018), they did not account for genome-wide heterogeneity in introgression among strains during hybridization. Much rather, analyses of ancestry disequilibrium suggest a role for negative epistasis between multiple ancestry-specific loci spread across the genome rather than single major effect mutations such as selfish elements or meiotic drivers (Zanders et al., 2014; Hu et al., 2017; Nuckolls et al., 2017). Functional work is needed to identify the genetic elements conveying reproductive isolation.

## Material and Methods

### Strains

This study is based on a global collection of *S. pombe* consisting of 161 world-wide distributed strains (see **Supplementary Table 1**) described in Jeffares *et al*. (2015).

### Inferring ancestry components

To characterize genetic variation across all strains, we made use of publically available data in variant call format (VCF) derived for all strains from Illumina sequencing with an average coverage of around 80x (Jeffares et al., 2015). The VCF file consists of 172,935 SNPs obtained after read mapping to the *S. pombe* 972 h^−^ reference genome (ASM294v264) (Wood et al., 2003) and quality filtering (see **Supplementary Table 1** for additional information). We used a custom script in R 3.4.3 (Team, 2014) with the packages *gdsfmt* 1.14.1 and *SNPRelate* 1.12.2 (Zheng et al., 2017, 2012), to divide the VCF file into genomic windows of 200 SNPs with overlap of 100 SNPs. This resulted in 1925 genomic windows of 1 - 89 kb in length (mean 13 kb). For each window, we performed principal component analyses (PCA) using *SNPRelate* 1.12.2 (Zheng et al., 2017, 2012) (example in **Figure 1a** and **Supplementary Figure 1**). The proportion of variance explained by the major axis of variation (PC1) was consistently high and allowed separating strains into two genetic groups/clusters, *Sp* and *Sk* (see main text, **Figure 1b**). We calculated population genetic parameters within clusters including pairwise nucleotide diversity (*π*) (Nei and Li, 1979), Watterson theta (*θ_w_*) (Watterson, 1975), and Tajima’s *D* (Tajima, 1989), as well as the average number of pairwise differences between clusters (*D_xy_*) (Nei and Li, 1979) using custom scripts. Statistical significance of the difference in nucleotide diversity (*π*) between ancestral clades was inferred using Kendall’s τ as test statistic. Since values of adjacent windows are statistically non-independent due to linkage, we randomly subsampled 200 windows along the genome with replacement. This was repeated a total of 10 times for each test statistic, and we report the maximum p-value. Given the consistent difference between clusters (**Figure 1** and **Supplementary Figure 2, 3 and 5**), normalised PC score could be used to attribute either *Sp* (low-diversity) or *Sk* (high-diversity) ancestry to each window (summary statistics for each window are given in **Supplementary Table 2**). This was performed both for the subset of 57 samples (**Figure 1c**) and for all 161 samples (**Supplementary Figure 6**). Using different window sizes (100, 50 and 40 SNPs with overlap of 50, 25 and 20 respectively) yielded qualitatively the same results. Intermediate values in PC1 (between 0.25 and 0.75) were only observed in few, sequential windows where samples transitioned between clusters (**Supplementary Figure 3**). The only exception was sample JB1207, which we found to be diploid (for details see below).

### Population structure after hybridization

To characterise the genome-wide distribution of ancestry components along the genome, we ran a hierarchical cluster analysis on the matrix containing ancestry information (*Sp* or *Sk*) for each window (columns) and strain (rows) using the R package *Pvclust 2.0.0* (Suzuki and Shimodaira, 2006). *Pvclust* includes a multiscale bootstrap resampling approach to calculate approximately unbiased probability values (p-values) for each cluster. We specified 1000 bootstraps using the Ward method and a Euclidian-based dissimilarity matrix. The analysis was run both for the whole genome (**Figure 1c**) and by chromosome (**Figure 1c**, **Supplementary Figure 7**).

### Phylogenetic analysis of the mitochondrial genome

From the VCF file, we extracted mitochondrial variants for all 161 samples (Jeffares et al., 2015) and generated an alignment in *.fasta* format by substituting SNPs into the reference *S. pombe* 972 h–reference genome (ASM294v264) using the package *vcf2fasta* (https://github.com/JoseBlanca/vcf2fasta/, version Nov. 2015). We excluded variants in mitochondrial regions with SVs inferred from long reads. A maximum likelihood tree was calculated using *RaxML* (version 8.2.10-gcc-mpi) (Stamatakis, 2014) with default parameters, GTRGAMMAI approximation, final optimization with GTR + GAMMA + I and 1000 bootstraps. The final tree was visualised using *FigTree* 1.4.3 (http://tree.bio.ed.ac.uk/software/figtree/) (**Supplementary Figure 8**).

### Time of hybridization

Previous work (Jeffares et al., 2015) has shown that the time to the most recent common ancestor for 161 samples dates back to around 2300 years ago. This defines the maximum boundary for the time of hybridization. We used the theoretical model by Janzen et al., (2018) to infer the age of the initial hybridization event. The model predicts the number of ancestry blocks and junctions present in a hybrid individual as a function of time and effective population size (*N_e_*). First, we obtained an estimate of *N_e_* using the multiple sequential Markovian coalescent (MSMC). We constructed artificial diploid genomes from strains with consistent clustering by ancestry (**Figure 1c**) and estimated change in *N_e_* as function across time using *MSMC* 2-2.0.0 (Schiffels and Durbin, 2014). In total we took four samples per group and produced diploid genomes in all possible six pairs for each group, except for one cluster that had only two samples (JB1205 and JB1206). Bootstraps were produced for each analysis, subsampling 25 genomic fragments per chromosome of 200 kb each. Resulting effective population size and time was scaled using reported mutation rate of 2 · 10^−10^ mutations site^-1^ generation^-1^ (Farlow et al., 2015). Although it is difficult to be certain of the number of independent hybridization events, it is interesting to see that some clusters show similar demographic histories (**Supplementary Figure 21**). Regardless of the demographic history in each cluster, long-term *N_e_* as estimated by the harmonic mean ranged between 1 · 10^5^ and 1 · 10^9^. *N_e_* of the near-pure ancestral *Sp* and *Sk* cluster was 7 · 10^5^ and 9 · 10^6^, respectively. These estimates of *N_e_* are consistent with previous reports of 1 · 10^7^ (Farlow et al., 2015).

We then used a customised R script with the ancestral component matrix to estimate the number of ancestry blocks (*Sp* or *Sk* clade) (**Supplementary Figure 10**). We used the R script from Janzen et al., (2018), and ran the model in each sample and chromosome using: *N_e_* = 1 · 10^6^, *r* = number of genomic windows per chromosome, *h0* = 0.298 (mean heterogenicity (h0) was estimated from the ancestral haplotype matrix) and *c* = 7.1, 5.6, and 4.1 respectively for chromosome I, II and III (values taken from Munz et al. (1989)) (**Supplementary Figure 11**). Given the large *N_e_*, no changes in mean heterogenicity is expected over time after hybridization due to drift (the proportion of ancestral haplotypes *Sp* and *Sk* in hybrids, estimated as 2*pq*, where *p* and *q* are the proportion of each ancestral clade in hybrids). Accordingly, results did not change within the range of the large *N_e_* values. For this analysis, samples with proportion of admixture lower than 0.1 were excluded.

### Phenotypic variation and reproductive isolation

We sourced phenotypic data of 229 phenotypic measurements in the 161 strains including amino acid quantification on liquid chromatography (aaconc), growth and stress on solid media (smgrowth), cell growth parameters and kinetics in liquid media (lmgrowth) and cell morphology (shape1 and shape2) from Jeffares et al. (2015). Data on reproductive isolation measured as the percentage of viable spores in pairs of crosses were compiled from Jeffares et al. (2015) and Marsellach (2017). A summary of all phenotypic measurements and reproductive data is provided in **Supplementary Table 4** and **5**, respectively.

First, we normalized each phenotypic trait y using rank-based transformation with the relationship normal.y = qnorm(rank (y) / (1 + length(y))). We then conducted PCA on normalized values of all phentoypic traits using the R package *missMDA* 1.12 (Josse and Husson, 2016). We estimated the number of dimensions for the principal component analysis by cross-validation, testing up to 30 PC components and imputing missing values. In addition to PCA decomposing variance across all traits, we examined the effect of admixture on each trait separately. Samples were divided into three discrete categories of admixture: two groups including samples with low admixture proportions (proportion of *Sp* or *Sk* clades higher than 0.9), and one for hybrid samples (proportion of *Sp* or *Sk* clades between 0.1 to 0.9).

Significant differences in phenotypic distributions between groups were tested using *Tukey Honest Significant Differences* as implemented in *Stats 3.4.2* (Team, 2014). **Supplementary Figure 12** shows the distribution of phenotypic values by admixture category for each trait. The number of traits with significant differences among groups was contrasted to values obtained by randomising admixture categories without replacement (permutations of the *Sp, Sk*, or *hybrid* category). Observed values were contrasted with distribution of the expected number of significant traits after running 10000 independent permutations (**Supplementary Figure 13**).

### Heritability

Heritability was estimated for all normalized traits using *LDAK* 5.94 (Speed et al., 2012), calculating independent kinship matrices derived from: 1) all SNPs and 2) ancestral haplotypes. Both SNPs and haplotype data were binary encoded (0 or 1). Jeffares et al. (2015) showed that heritability estimates between normalised and raw values are highly correlated (*r* = 0.69, *p*-value ≤ 2.2 · 10^−16^). Heritability estimated with SNP values were strongly correlated with those from ancestral haplotypes (*r* = 0.82, *p*-value ≤ 2.2 · 10^−16^). Heritability estimates and standard deviation for each trait for both SNP and ancestral haplotypes are detailed in **Supplementary Table 6**.

### Identification of ploidy changes

*S. pombe* is generally considered haploid under natural conditions. Yet, for two samples ancestry components did not separate on the principle component axis 1 (see above) for much of the genome. Instead, these samples were intermediate in PC1 score. A possible explanation is diploidisation of the two ancestral genomes. To establish the potential ploidy of samples, we called variants for all 161 samples using the Illumina data from Jeffares *et al* (2015). Cleaned reads were mapped with *BWA* (version 0.7.17-r1188) in default settings and variants were called using *samtools* and *bcftools* (version 1.8). After filtering reads with a QUAL score > 25, the number of heterozygous sites per base per 20kb window were calculated. Additionally the nuclear content (C) as measured by Jeffares et al. (2015) (**Supplementary Table S4** in Jeffares *et al* (2015)) were used to verify increased ploidy. Two samples showed high heterozygosity along the genome (JB1169 and JB1207) of which JB1207 for which data were available also showed a high C-value, suggesting that these samples are diploid (**Supplementary Figures 4 & 22**). In JB1207, heterozygosity varies along the genome, with regions of high and low diversity. Assigning ancestry (see **Supplementary Figure 6**), shows that the haploid parents differed from each other and that both chromosomes stem from hybrids between the *Sp* and *Sk* clades. Sample JB1110 showed genomic content similar to JB1207, but did not show heterozygosity levels above that of haploid strains, suggesting the increase in genome content occurred by autoploidization.

### High-weight genomic DNA extraction and whole genome sequencing

To obtain high weight gDNA for long-read sequencing, we grew strains from single colonies and cultured them in 200 mL liquid EMM at 32 °C shaking at 150 r.p.m. overnight. Standard media and growth conditions were used throughout this work (Hagan et al., 2016) with minor modifications: We used standard liquid Edinburgh Minimal Medium (EMM; Per liter: Potassium Hydrogen Phthalate 3.0 g, Na HPO_4_·2H_2_O 2.76 g, NH_4_Cl 5.0 g, D-glucose 20 g, MgCl_2_·6H_2_O 1.05 g, CaCl_2_·2H_2_O 14.7 mg, KCl 1 g, Na_2_SO_4_ 40 mg, Vitamin Stock ×1000 1.0 ml, Mineral Stock ×10,000 0.1 ml, supplemented with 100 mg 1^−1^ adenine and 225 mg 1^−1^ leucine) for the asexual growth. DNA extraction was performed with Genomic Tip 500/G or 100/G kits (Qiagen) following the manufacturer’s instruction, but using Lallzyme MMX for lysis (Flor-Parra et al 2014, doi:10.1002/yea.2994). For each sample, 20 kb libraries were produced that were sequenced on one SMRT cell per library using the Pacific Biosciences RSII Technology Platform (PacBio^®^, CA). For a subset of eight samples, additional sequencing was performed using Oxford Nanopore (MinION). Sequencing was performed at SciLifeLab, Uppsala, Gene centre LMU, Munich and The Genomics & Bioinformatics Laboratory, University of York. We obtained on average 80x (SMRT) and 140x (nanopore) coverage for the nuclear genome for each sample (summary in **Supplementary Table 7**).

Additionally, 2.5 μg of the same DNA was delivered to the SNP&SEQ Technology Platform at the Uppsala Biomedical Centre (BMC), for Illumina sequencing. Libraries were prepared using the TruSeq PCRfree DNA library preparation kit (Illumina Inc.). Sequencing was performed on all samples pooled into a single lane, with cluster generation and 150 cycles paired-end sequencing on the HiSeqX system with v2.5 sequencing chemistry (Illumina Inc.). These data were used for draft genome polishing (see below).

### *De novo* assembly of single-molecule read data

*De novo* genomes were assembled with *Canu* 1.5 (Koren et al., 2017) using default parameters. BridgeMapper from the *SMRT* 2.3.0 package was used to polish and subsequently assess the quality of genome assembly. Draft genomes were additionally polishing using short Illumina reads, running four rounds of read mapping to the draft genome with *BWA* 0.7.15 and polishing with *Pilon* 1.22 (Walker et al., 2014). Summary statistics of the final assembled genomes are found in **Supplementary Table 7**. *De novo* genomes were aligned to the reference genome using *MUMmer* 3.23 (Kurtz et al., 2004). Contigs were classified by reference chromosome to which they showed the highest degree of complementary. We used customised python scripts to identify and trim mitochondrial genomes.

### Structural variant detection

Structural variants (SVs) were identified by a combination of a *de novo* and mapping approach. *De novo* genomes were aligned to the reference genome using *MUMmer*, and SVs were called using the function show-diff and the package *SVMU* 0.2beta (Khost et al., 2016). Then, raw long reads were mapped to the reference genome with *NGMLR* and genotypes were called using the package *Sniffles* (Sedlazeck et al., 2018). We implemented a new function within Sniffles “forced genotypes”, which calls SVs by validating the mapping calls from an existing list of breaking points or SVs. This reports the read support per variant even down to a single read. We forced genotypes using the list of *de novo* breaking points to generate a multi-sample VCF file. SVs were merged using the package *SURVIVOR* (Jeffares et al., 2017) option merge with a threshold of 1kbp and requiring the same type. In total, it resulted in a list of 1498 SVs with 892 in common between the mapping and *de-novo* approaches (**Supplementary Figure 14**).

Within the 892 common variants we compared the accuracy of genotyping between sample by comparing genotypes obtained from *de novo* genomes and by mapping reads to reference genome. Additionally, we compared genotypes in samples sequenced with both PacBio and MinIon. In total we sequenced 8 samples with both technologies. We found high consistency for variants called with both sequencing technologies and observed that allele frequencies were highly correlated (r = 0.98, p-value ≤ 2.2 × 10^−16^) (**Supplementary Figures 14 - 17**). Only common SVs between the mapping and *de-novo* approach were considered, and variants with consistency below 50% were removed. We manually checked large SVs (larger than 10kb) by comparing the list of SVs with the alignment of the *de novo* genomes to the reference genome from MUMmer. This resulted in a final data set with 832 SVs (**Supplementary Table 8**).

### Distribution of SVs around developmentally programmed DNA double-strand breaks (DSB)

We tested the association between DSB and SVs by comparing the physical genomic coordinates of the final list of SV with DSB locations accessed from Fowler et al., (2014). Maintaining the same number of SV per chromosome, we used a customized R script to randomise SV coordinates and measure the distances to the closest DSB. We counted the number of SV present within different intervals of physical genetic distance ([0,500), [500, 1000), [1000, 2000), [2000, 4000), [4000, 10000), [10000, 20000), [20000, 30000) bp). Empirically observed values were contrasted with randomized distribution after running 10000 independent permutations. P-values of differences between randomization and observed values were obtained from the fraction of expected values higher than the observed value from the original data (**Supplementary Figures 18**).

### PCR validation of large SVs

To test the frequency of large inversions and rearrangements observed from long read data, we performed PCR verification over the breakpoints in the 57 non-clone samples. PCR was performed for both sides of the breakpoints, with a combination of one primer ‘outside’ of the inversion and both primers ‘inside’ the inversion (**Supplementary Figure 23**). PCR were performed on DNA using standard *Taq* polymerase, with annealing temperature at 59°C. The primers used, the coordinates in the reference and the expected amplicon length are given in **Supplementary Table 9**.

### Distribution of structural variants in ancestral population – Two dimensional folded site frequency spectrum

We used the location of break points of SVs to identify whether a variant was located in the *Sp* or *Sk* genetic background in each sample. Ancestral haplotypes are difficult to infer in telomeric and centromeric regions given the low confidence in SNP calling in those regions, resulting in low percentage of variance explained by PC1. Thus SVs with break points in those regions were excluded from this analysis (19 SVs). SVs were grouped by ancestral group and allele frequencies were calculated for each ancestral population. We used these frequencies to build a two dimensional folded site frequency spectrum (2dSFS). In order to compare this 2dSFS, we repeated the analysis using SNP data from all 57 samples. Considering that the majority of identified SVs with long reads were transposable elements, we also made use of LRT insertion-deletion polymorphism (indels) inferred from short reads. For this additional data we produced a similar folded 2d SFS. LTR indel data were taken from Jeffares et al., (2015) and are listed in **Supplementary Table 11**.

### Decay in linkage disequilibrium (LD)

To contrast LD between alleles from alternative ancestral groups, we calculated LD between all described genomic windows within chromosomes (**Supplementary Figure 20**). For this analysis only hybrid samples were considered (strains with admixture proportion higher than 0.1). For each pair of windows, we polarized windows by ancestry (at a threshold of > 0.7) and calculated standardized LD as the squared Pearson’s correlation coefficient (R^2^) (Hill and Robertson, 1968; Weir, 1979). This measurement takes into consideration difference in allele frequencies. The expected value of R^2^ (E(R^2^)) can be approximated by (Hill and Weir, 1988):

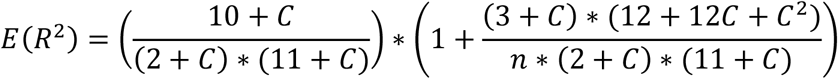

Where C corresponds to product between the genetic distance (bp) and the population recombination rate (ρ) in n number of haplotype sampled. The population recombination rate was calculated as: ρ = 4 * *N_e_* * *c*, where c is the recombination fraction between sites and Ne is the effective population size. We fitted a nonlinear model to obtain least squares estimates of ρ using a customized R script. The decay of LD with physical distance can be described with this model (Remington et al., 2001). LD values were grouped in three categories: i) comparison between windows with high proportion (*Sp*>0.7) of *Sp* ancestral group (*Sp-Sp*); ii) high proportion (*Sk*>0.7) of *Sk* ancestral group (*Sk-Sk*); and iii) high proportion of opposite ancestral groups (*Sp-Sk*). i) and ii) represent cases of positive ancestry disequilibrium, iii) will be denoted as negative ancestry disequilibrium.

### Data availability

Nanopore, single-molecule real time sequencing data and de-novo genomes are available at NCBI Sequence Read Archive, BioProject ID XXX.

## Acknowledgments

We thank Fidel Botero-Castro, Ana Catalán, Sebastian Höhna, Ulrich Knief, Claire Peart, Joshua Peñalba, Ricardo Pereira, Matthias Weissensteiner, (LMU Munich) and S. Lorena Ament-Velásquez (Uppsala University) for providing valuable intellectual input on the various analyses, sharing scripts and critically comment on the manuscript. We are further indebted to Bernadette Weissensteiner for extensive help with laboratory work and Saurabh Pophaly for bioinformatics support (LMU Munich). We further acknowledge support for data generation from the National Genomics Infrastructure, Uppsala, Sweden, the Gene Centre, Munich, Germany, Sally James and Peter Ashton from the Bioscience Technology Facility, Department of Biology, University of York, U.K, and James Chong, Department of Biology, University of York, U.K. The computational infrastructure was provided by the UPPMAX Next-Generation Sequencing Cluster and Storage (UPPNEX) project funded by the Knut and Alice Wallenberg Foundation and the Swedish National Infrastructure for Computing and the York Advanced Research Computing Cluster (YARCC), University of York, U.K. This study was funded by LMU Munich to JW and NHGRI UM1 HG008898 to FS.

## Contributions

ST, BN, DJ and JW conceived of the study; All analyses were performed by ST with contributions from FS in structural variation calling, JD in de novo assembly, BN in ancestral inference and population genetics parameters and DJ in phenotypic and heritability analyses; ST and JD assembled de-novo genomes; BN designed primers for PCR validation of structural variants; ST, BN, and JW wrote the manuscript with input from all other authors.

