## Supplementary Figures for "Ancestral admixture is the main determinant of global biodiversity in fission yeast"

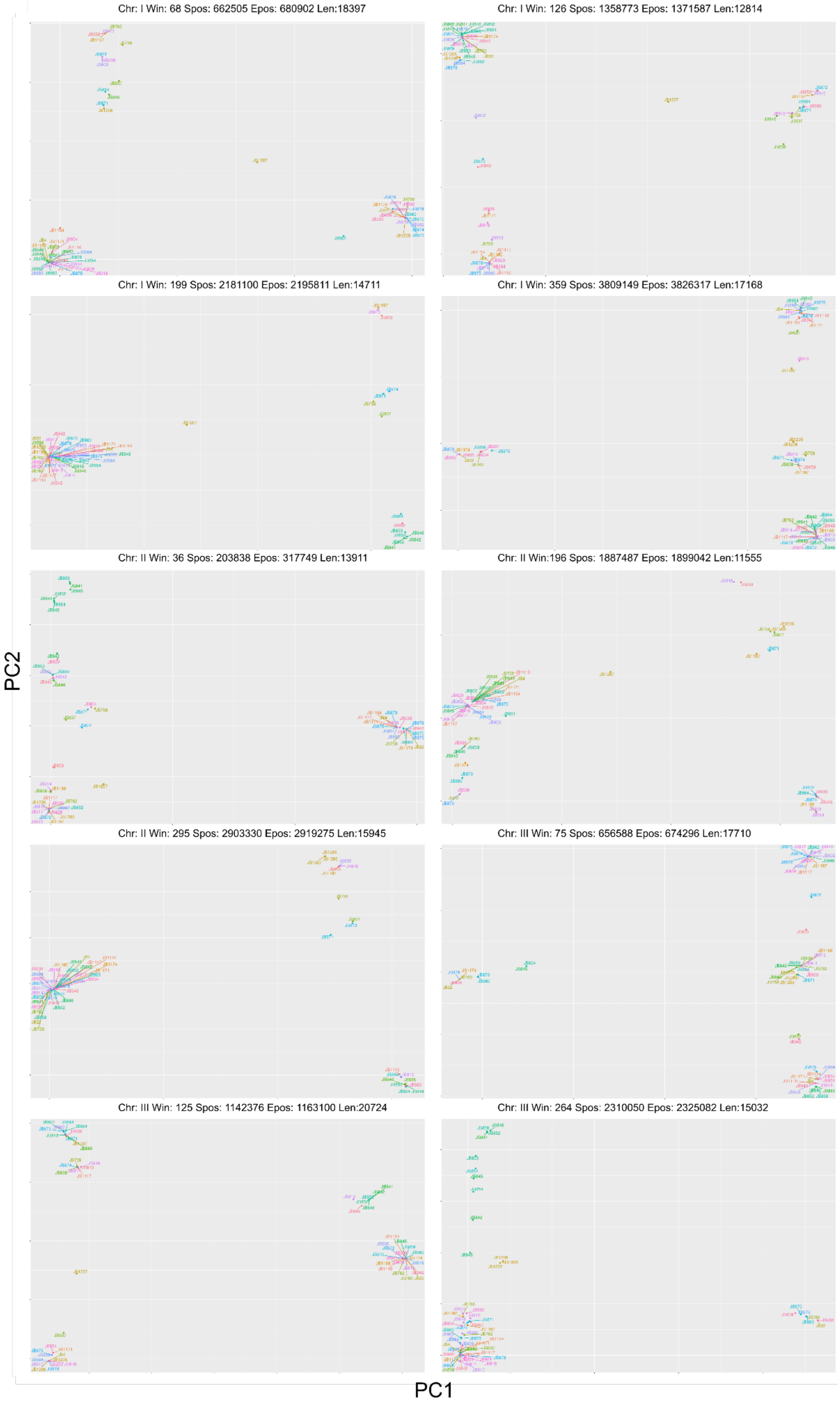
Chr: I Win: 68 Spos: 662505 Epos: 680902 Len:18397
Chr: I Win: 126 Spos: 1358773 Epos: 1371587 Len:12814
Chr: I Win: 199 Spos: 2181100 Epos: 2195811 Len:14711
Chr: I Win: 359 Spos: 3809149 Epos: 3826317 Len:17168
Chr: II Win: 36 Spos: 203838 Epos: 317749 Len:13911
Chr: II Win:196 Spos: 1887487 Epos: 1899042 Len:11555
Chr: II Win: 295 Spos: 2903330 Epos: 2919275 Len:15945
Chr: III Win: 75 Spos: 656588 Epos: 674296 Len:17710
Chr: III Win: 125 Spos: 1142376 Epos: 1163100 Len:20724
Chr: III Win: 264 Spos: 2310050 Epos: 2325082 Len:15032
PC2
PC1

**Supplementary Figure 1: Example of principal component analysis (PCA).** Each panel is an example PCA of different genomic windows. The chromosome (Chr.), window number (Win), start and end position (Spos and Epos) and length of the window in bp (Len) are shown in the header for each panel. Samples are differentiated with colours.

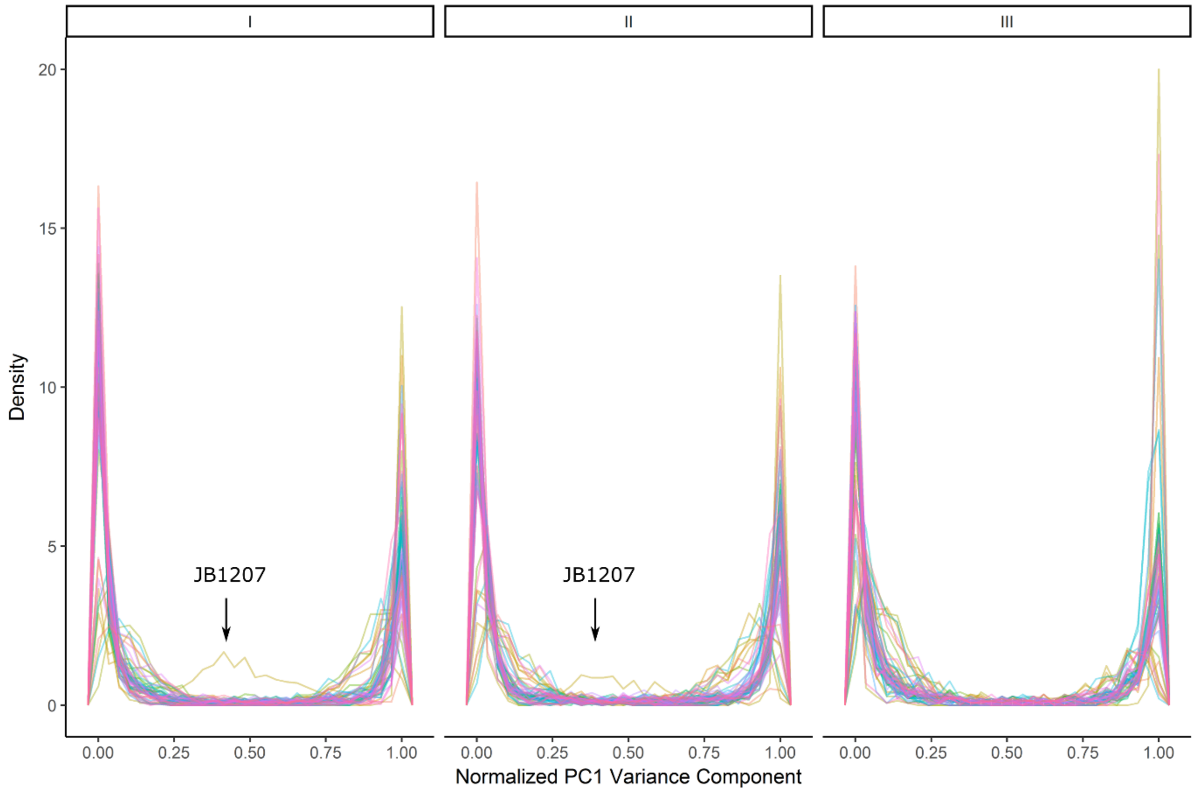


**Supplementary Figure 2: Normalised PC1 distribution.** Density plot of Normalised PC1 values, differentiating between chromosomes. Each line represent one sample. The sample JB1207 is highlighted as it is the only sample in high proportion of intermediate values.

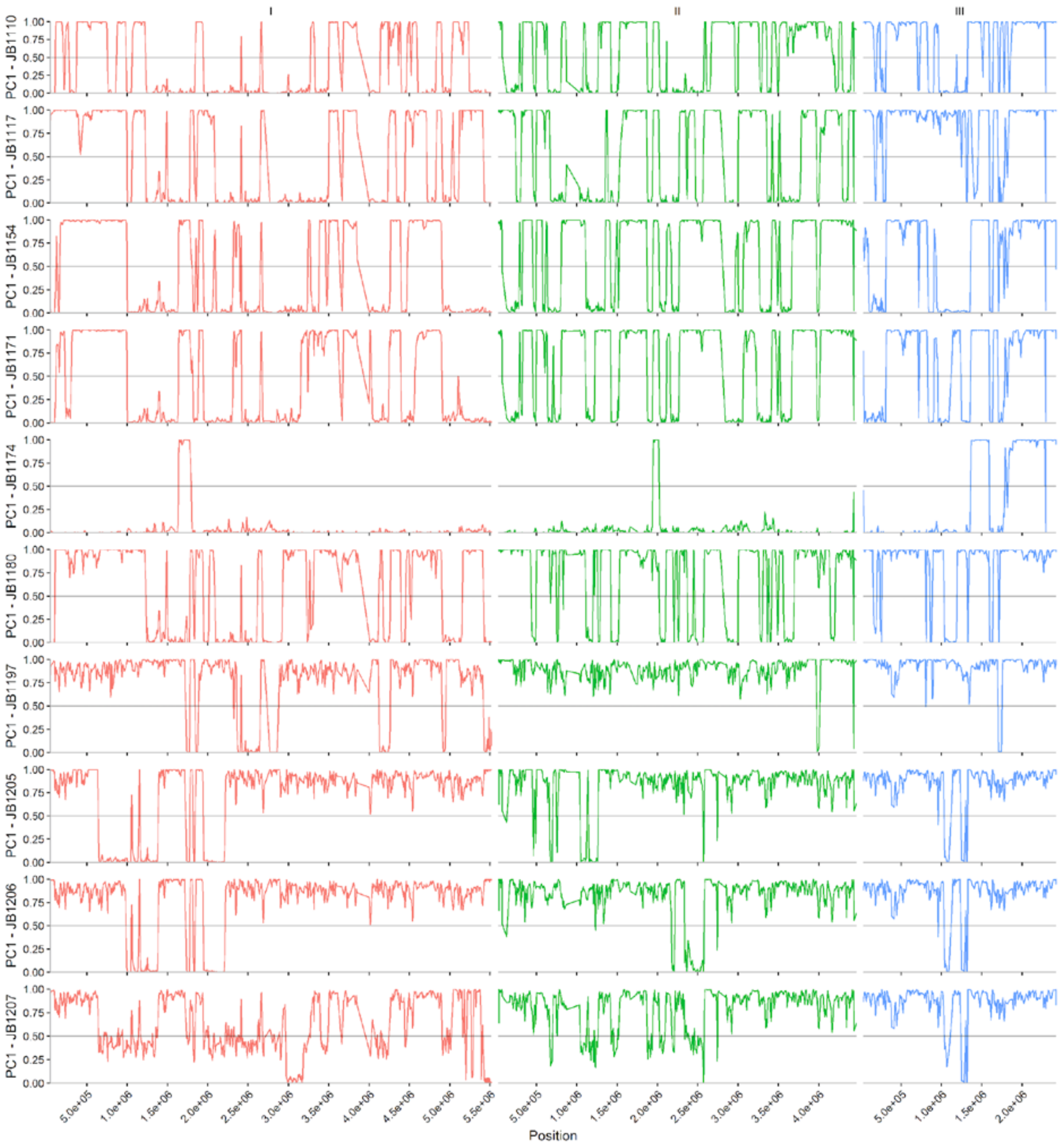


Figure continues on next page.

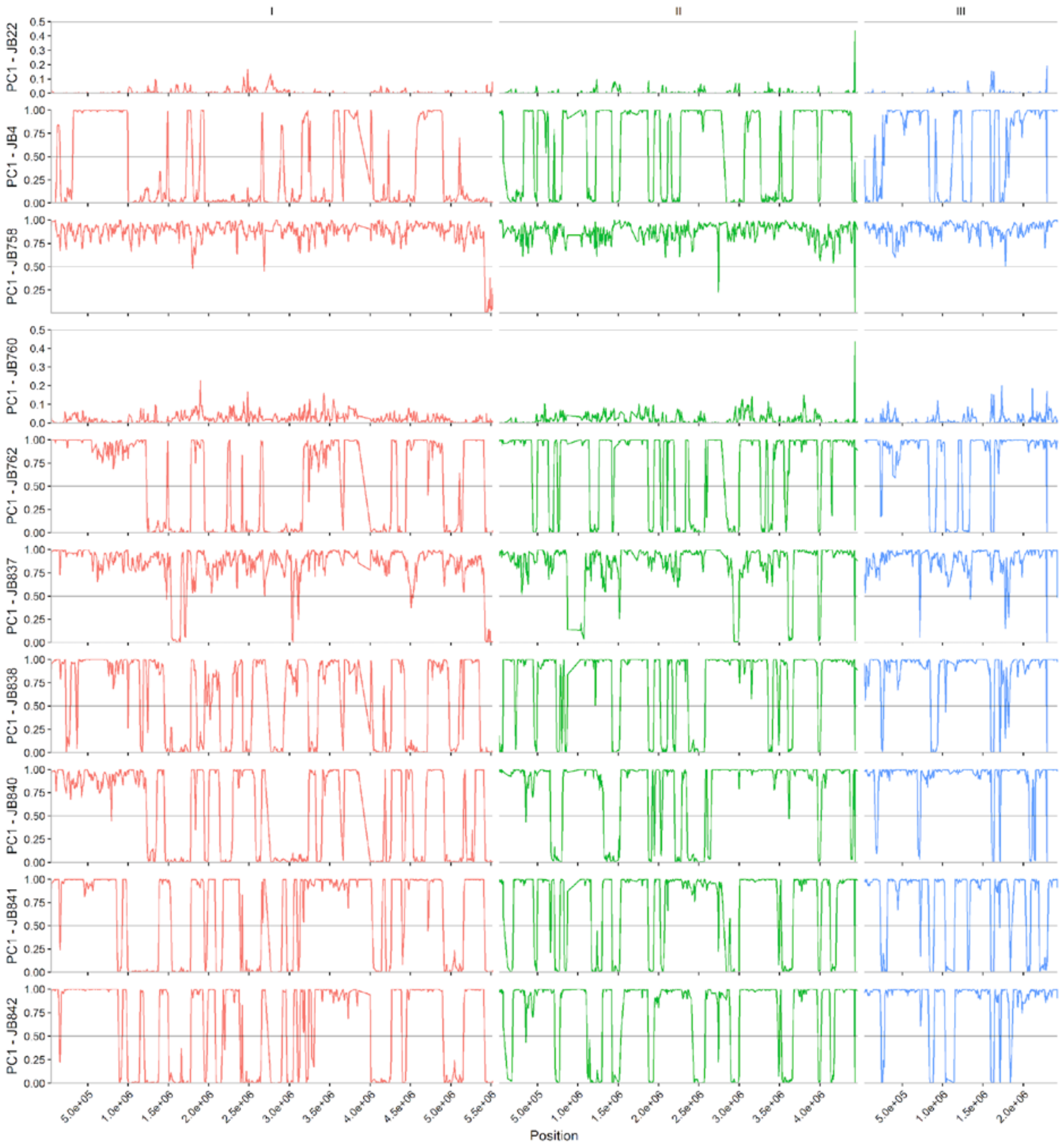


Figure continues on next page.

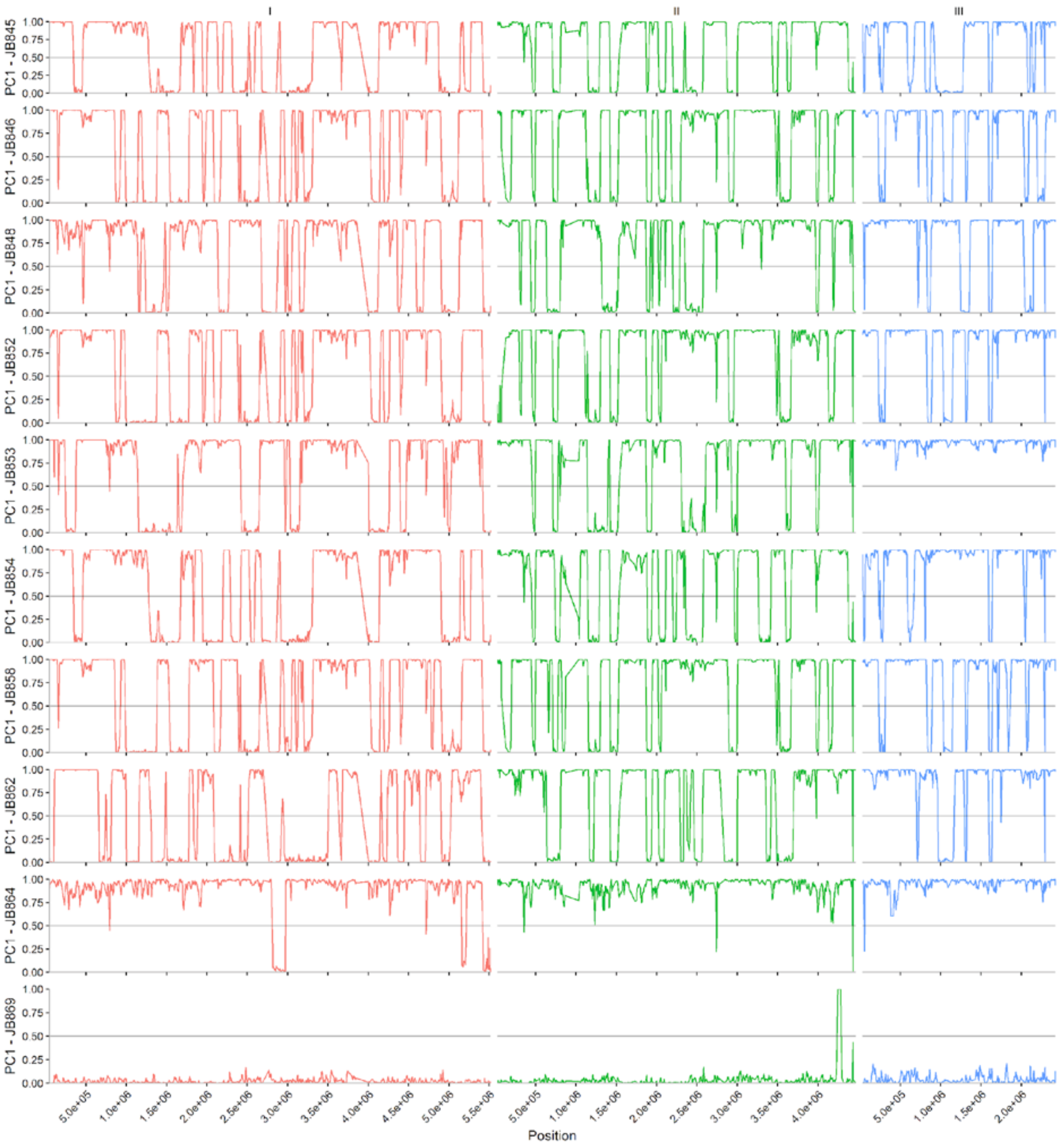


Figure continues on next page.

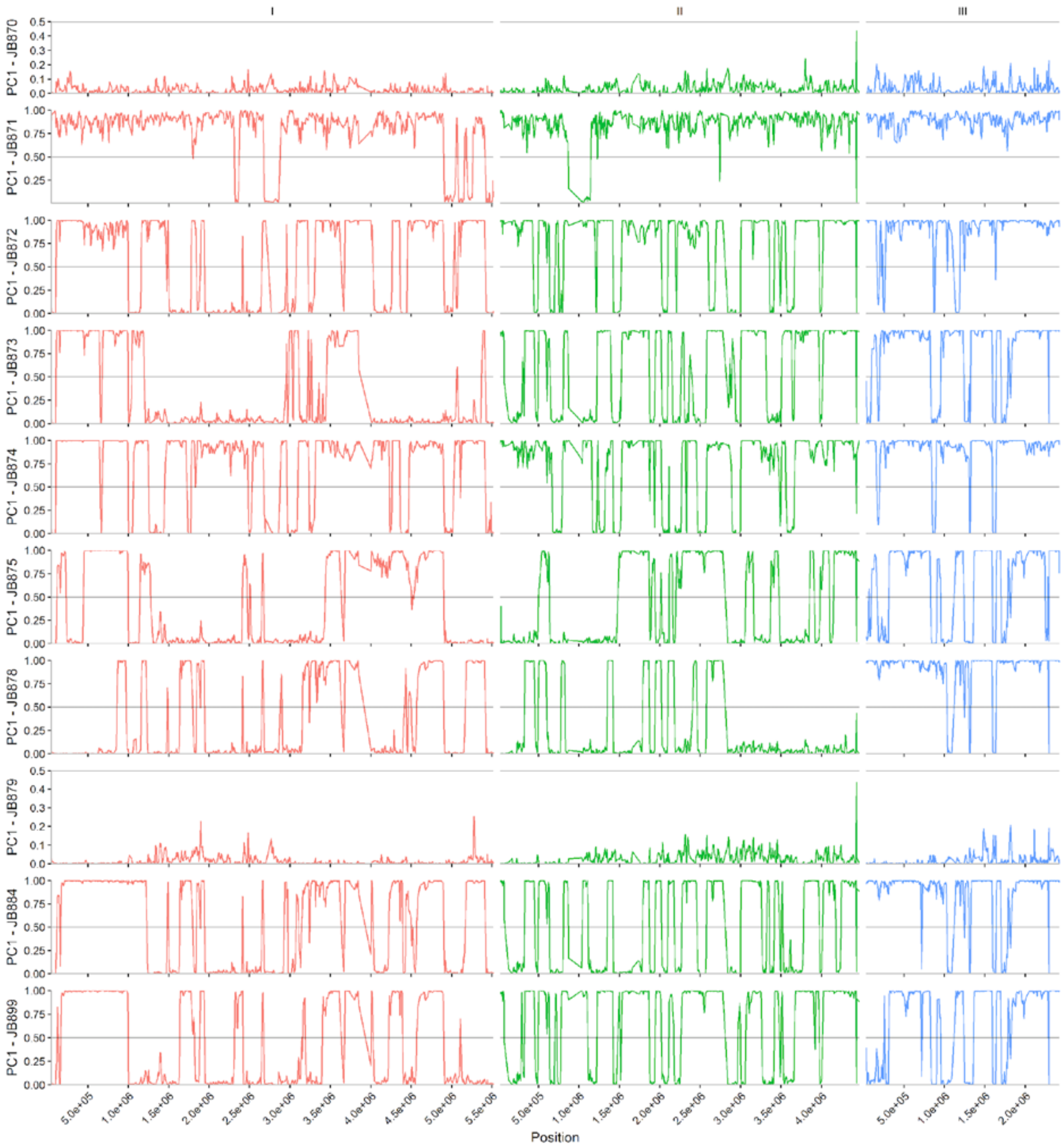


Figure continues on next page.

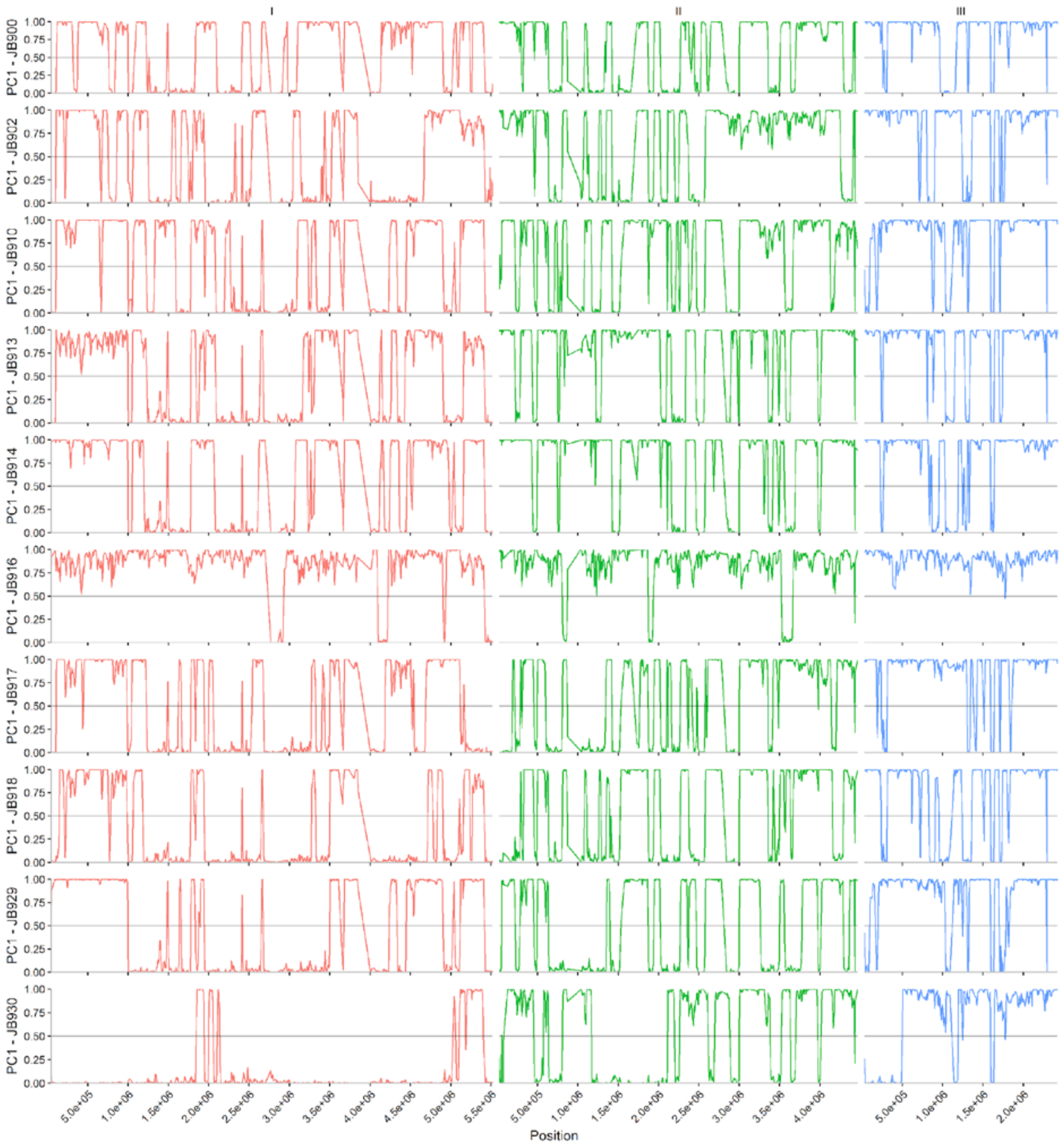


Figure continues on next page.

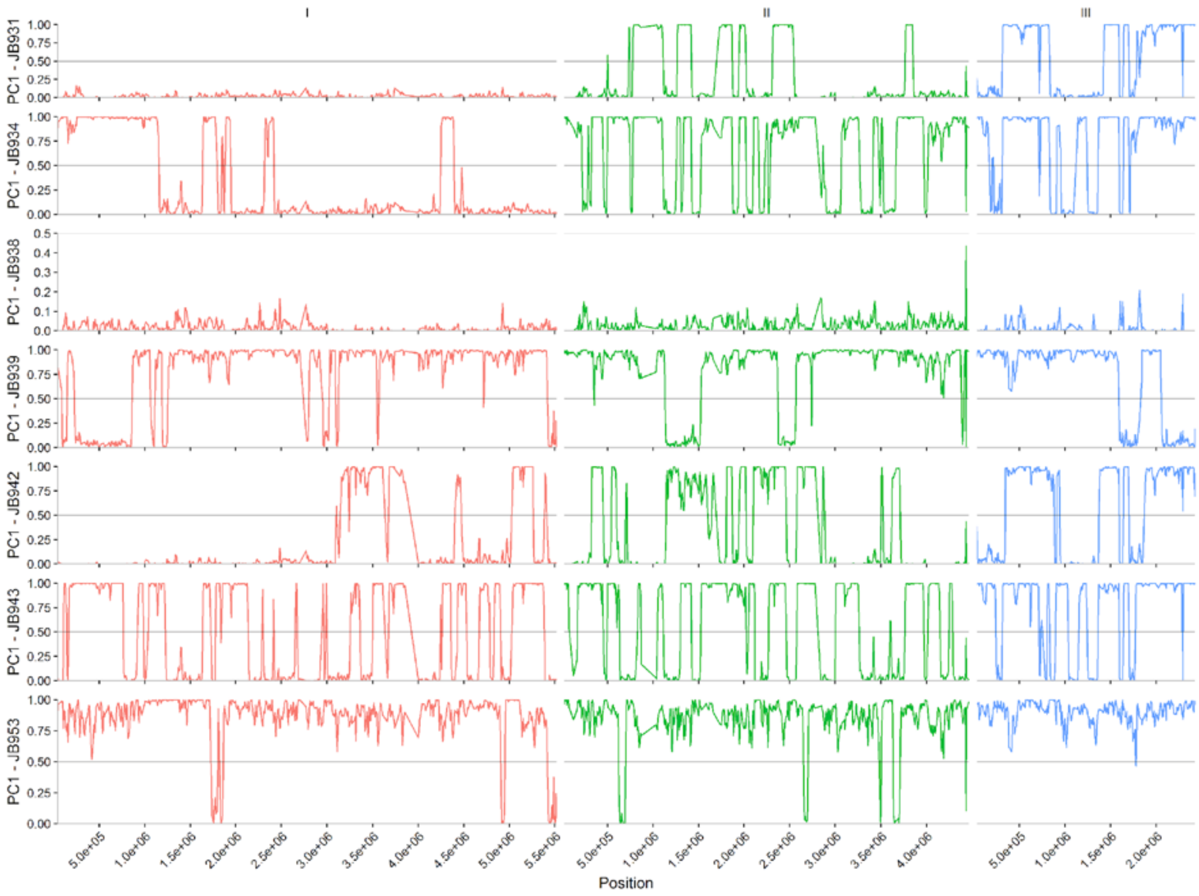


**Supplementary Figure 3: Distribution of Normalised PC1 values alone the genome.** Chromosomes are indicated by colour. Normalised PC1 values were polarized based on genetic diversity as described in Methods, values close to 0 or 1 represent genomic regions with the *Sp* or *Sk* haplotype, respectively.

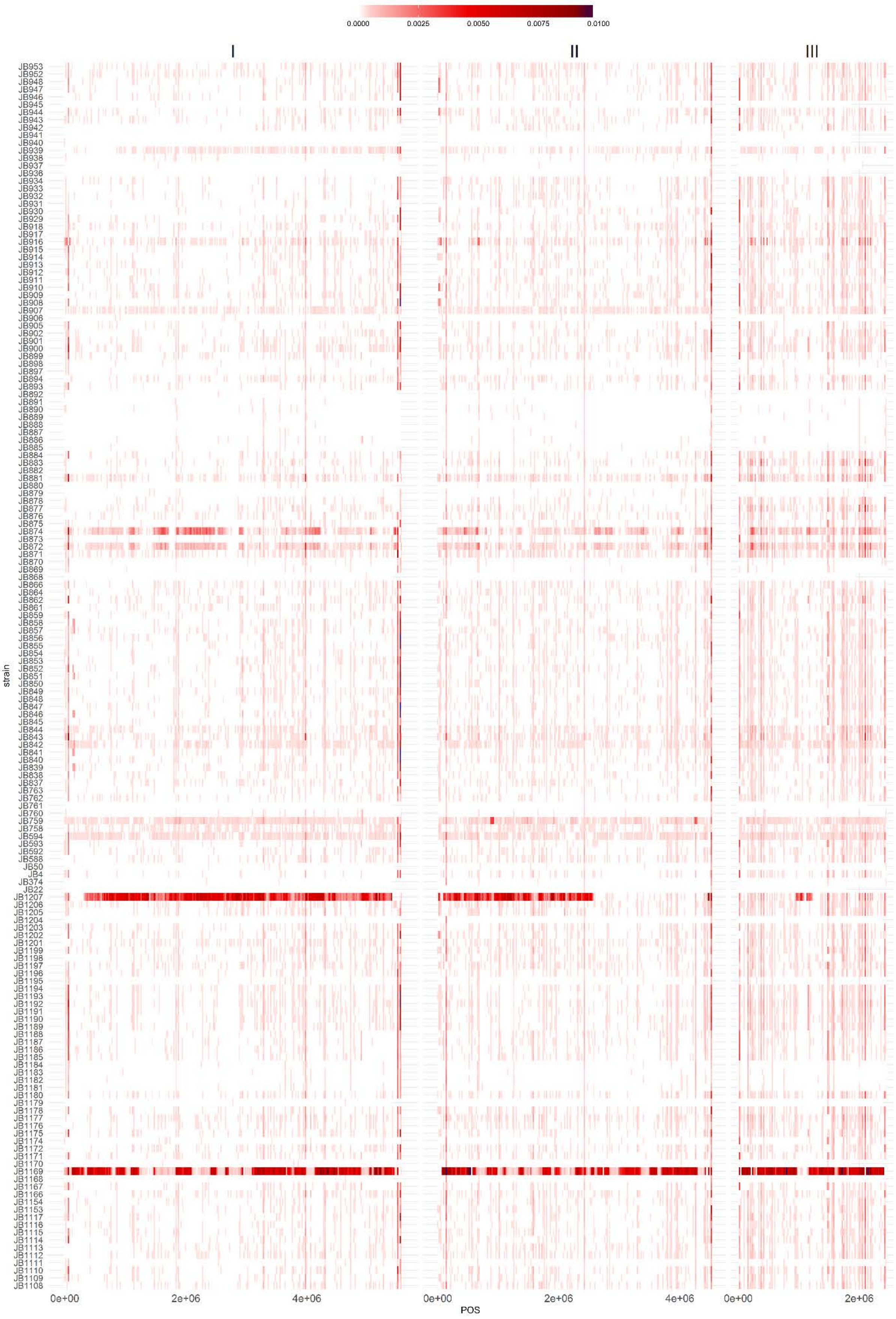


**Supplementary Figure 4: Heterozygosity along the genome.** Heatmap of heterozygosity per 20kb window and sample along the genome. High Heterozygosity values are shown in red. Strains JB1207 and JB1169 show high heterozygosity along large parts of the genome.

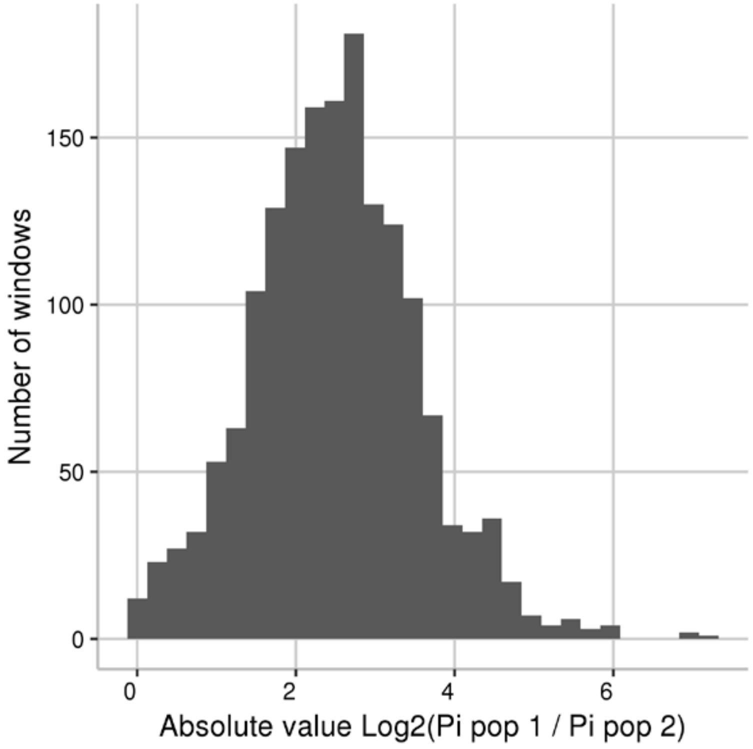


**Supplementary Figure 5: Difference in genetic diversity between ancestral groups.** Histogram of the ratio in genetic diversity ($\pi$) between the ancestral clade *Sk* over the *Sp* clade. Values in log 2 scale.

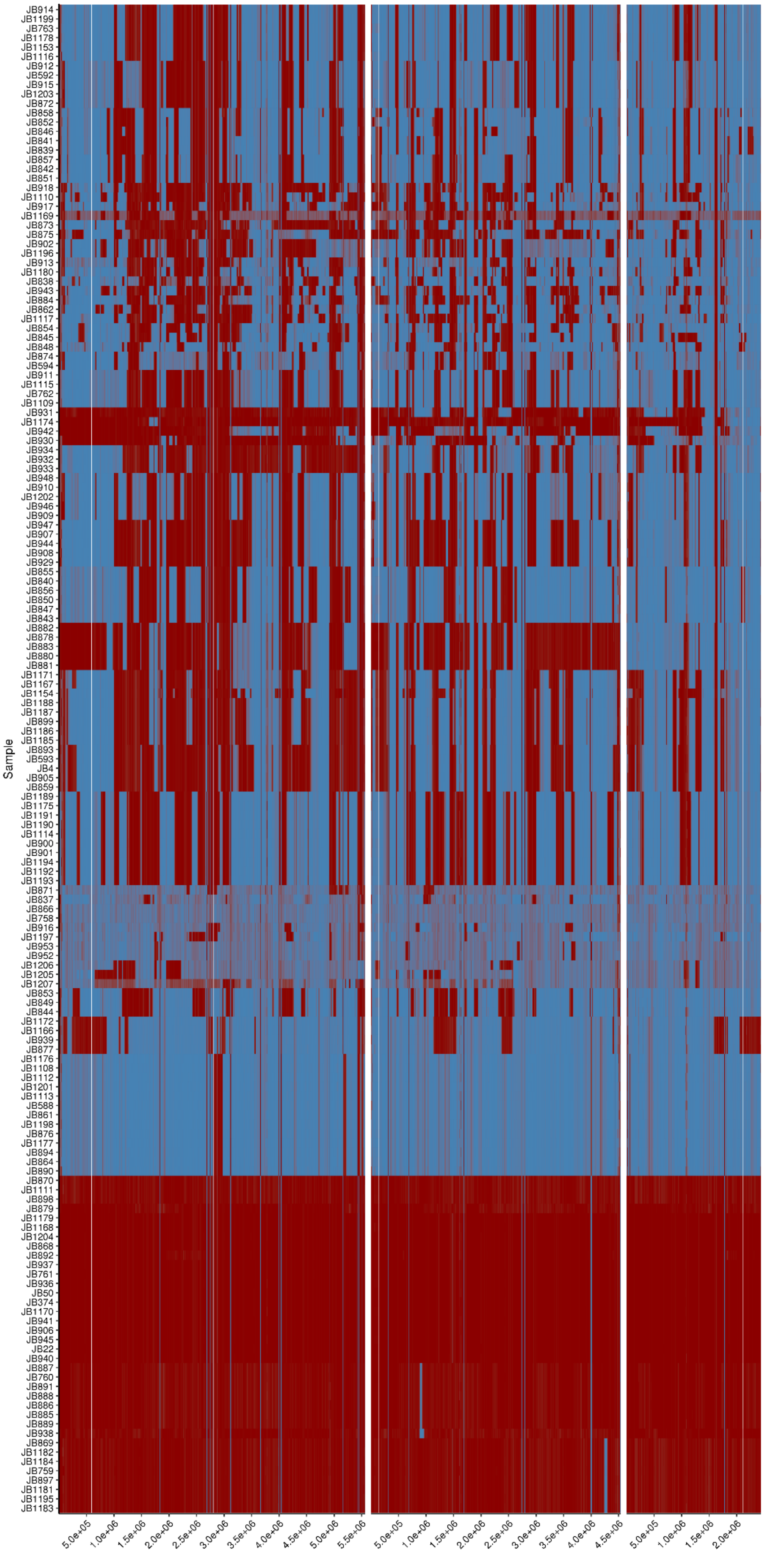

Sample
JB914
JB1199
JB763
JB1178
JB1153
JB1116
JB912
JB592
JB915
JB1203
JB872
JB858
JB852
JB846
JB841
JB839
JB857
JB842
JB851
JB918
JB1110
JB917
JB1169
JB873
JB875
JB902
JB1196
JB913
JB1180
JB838
JB943
JB884
JB862
JB1117
JB854
JB845
JB848
JB874
JB594
JB911
JB1115
JB762
JB1109
JB931
JB1174
JB942
JB930
JB934
JB932
JB933
JB948
JB910
JB1202
JB946
JB909
JB947
JB907
JB944
JB908
JB929
JB855
JB840
JB856
JB850
JB847
JB843
JB882
JB878
JB883
JB880
JB881
JB1171
JB1167
JB1154
JB1188
JB1187
JB899
JB1186
JB1185
JB893
JB593
JB4
JB905
JB859
JB1189
JB1175
JB1191
JB1190
JB1114
JB900
JB901
JB1194
JB1192
JB1193
JB871
JB837
JB866
JB758
JB916
JB1197
JB953
JB952
JB1206
JB1205
JB1207
JB853
JB849
JB844
JB1172
JB1166
JB939
JB877
JB1176
JB1108
JB1112
JB1201
JB1113
JB588
JB861
JB1198
JB876
JB1177
JB894
JB864
JB890
JB870
JB1111
JB898
JB879
JB1179
JB1168
JB1204
JB868
JB892
JB937
JB761
JB936
JB50
JB374
JB1170
JB941
JB906
JB945
JB22
JB940
JB887
JB760
JB891
JB888
JB886
JB885
JB889
JB938
JB869
JB1182
JB1184
JB759
JB897
JB1181
JB1195
JB1183
5.0e+05
1.0e+06
1.5e+06
2.0e+06
2.5e+06
3.0e+06
3.5e+06
4.0e+06
4.5e+06
5.0e+06
5.5e+06
5.0e+05
1.0e+06
1.5e+06
2.0e+06
2.5e+06
3.0e+06
3.5e+06
4.0e+06
4.5e+06
5.0e+05
1.0e+06
1.5e+06
2.0e+06

**Supplementary Figure 6: Distribution of ancestral *Sp* and *Sk* haplotypes along the genome.** Heatmap of ancestral *Sp* and *Sk* haplotypes along the genome for each sample. As in Figure 1c, but using all 161 samples.

(a)

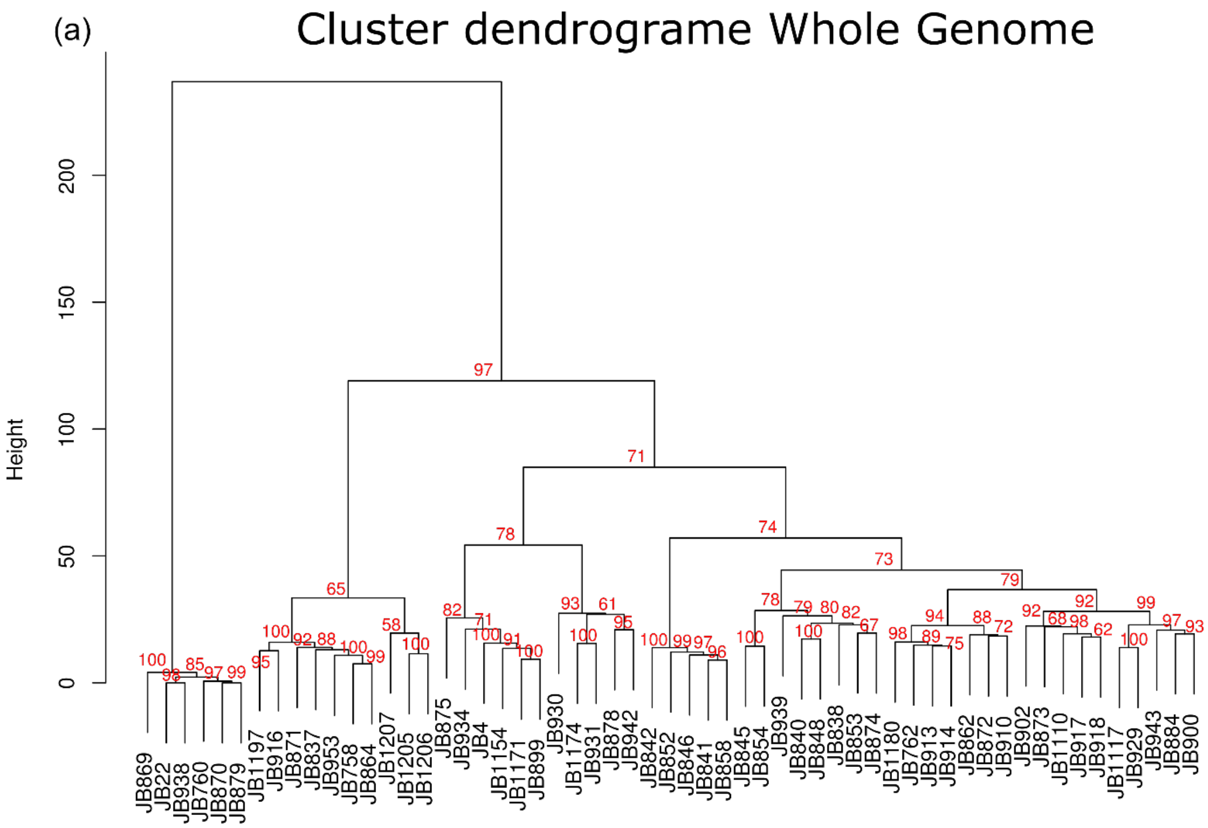


(b)

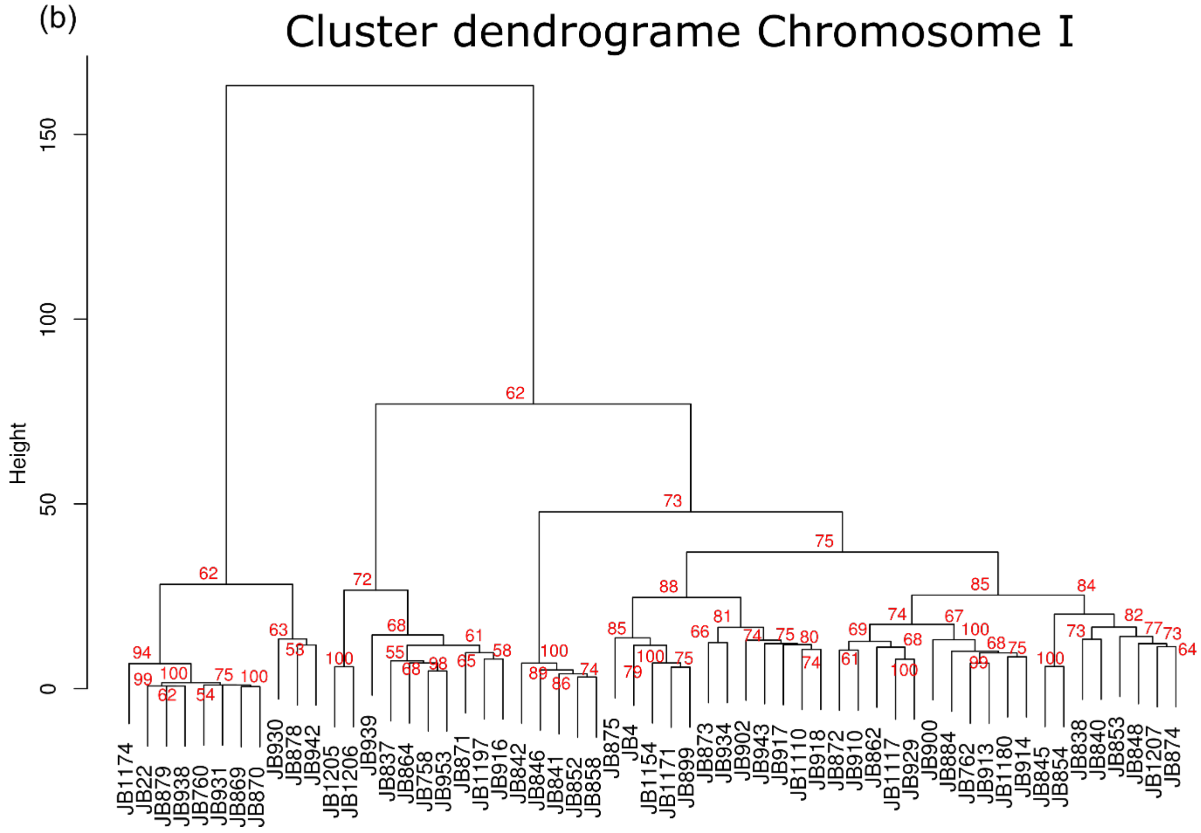


Figure continues on next page.

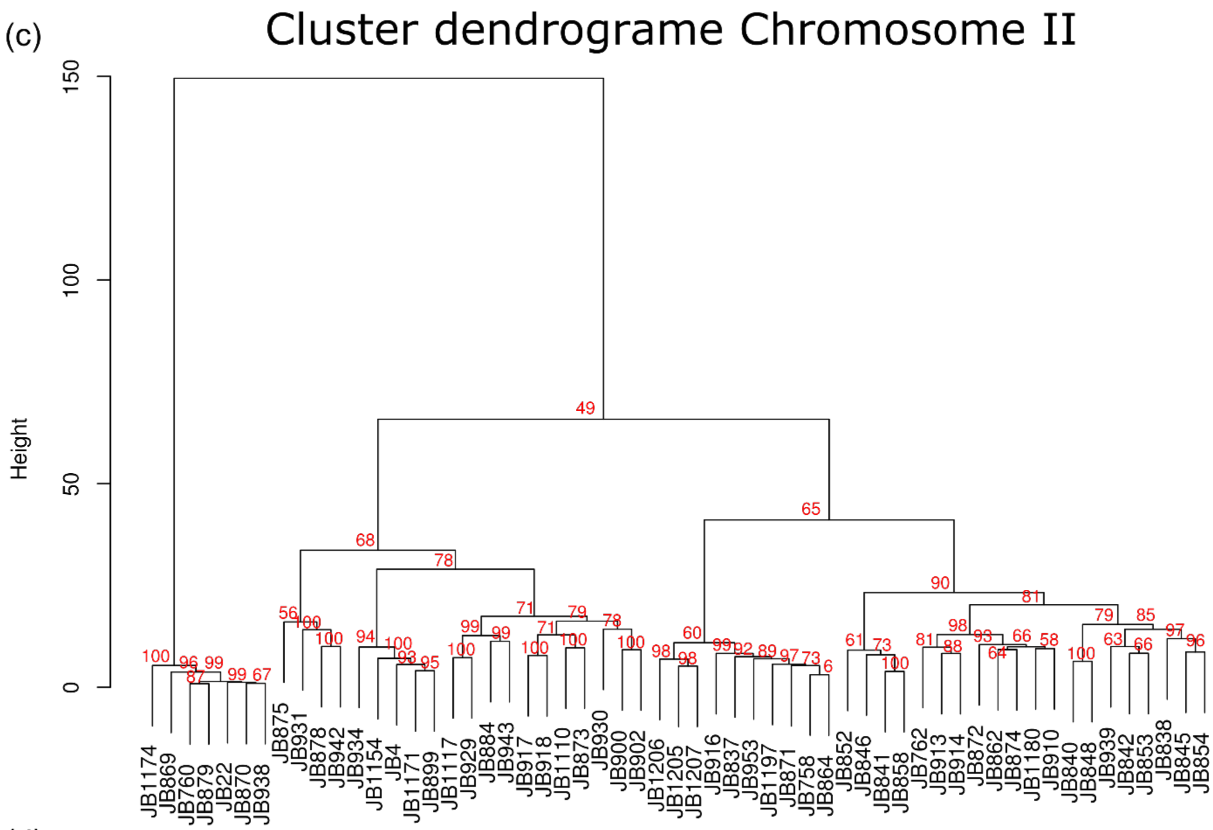


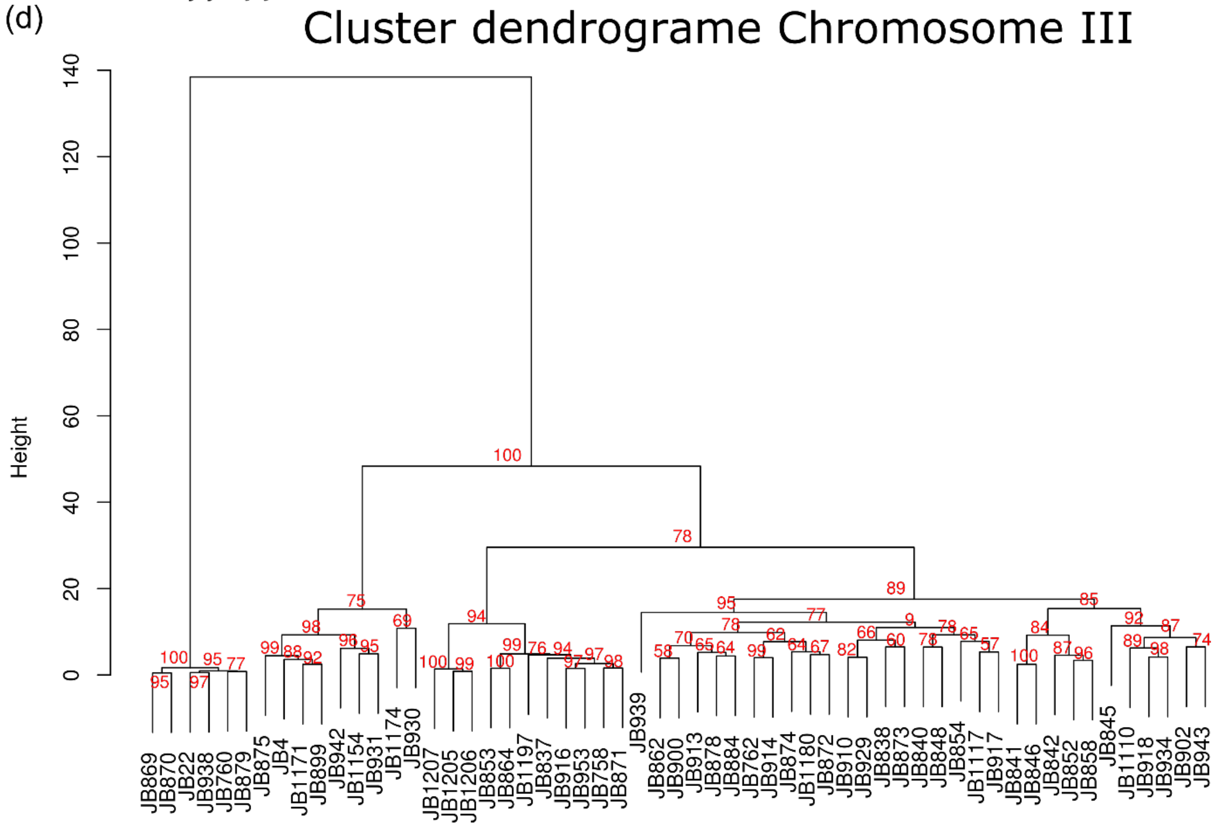


**Supplementary Figure 7: Clustering analyses by chromosome.** Clustering analyses using ancestral block distribution. Each panel shows the clustering using whole chromosome and independently by chromosome. Red numbers show bootstrapping support.

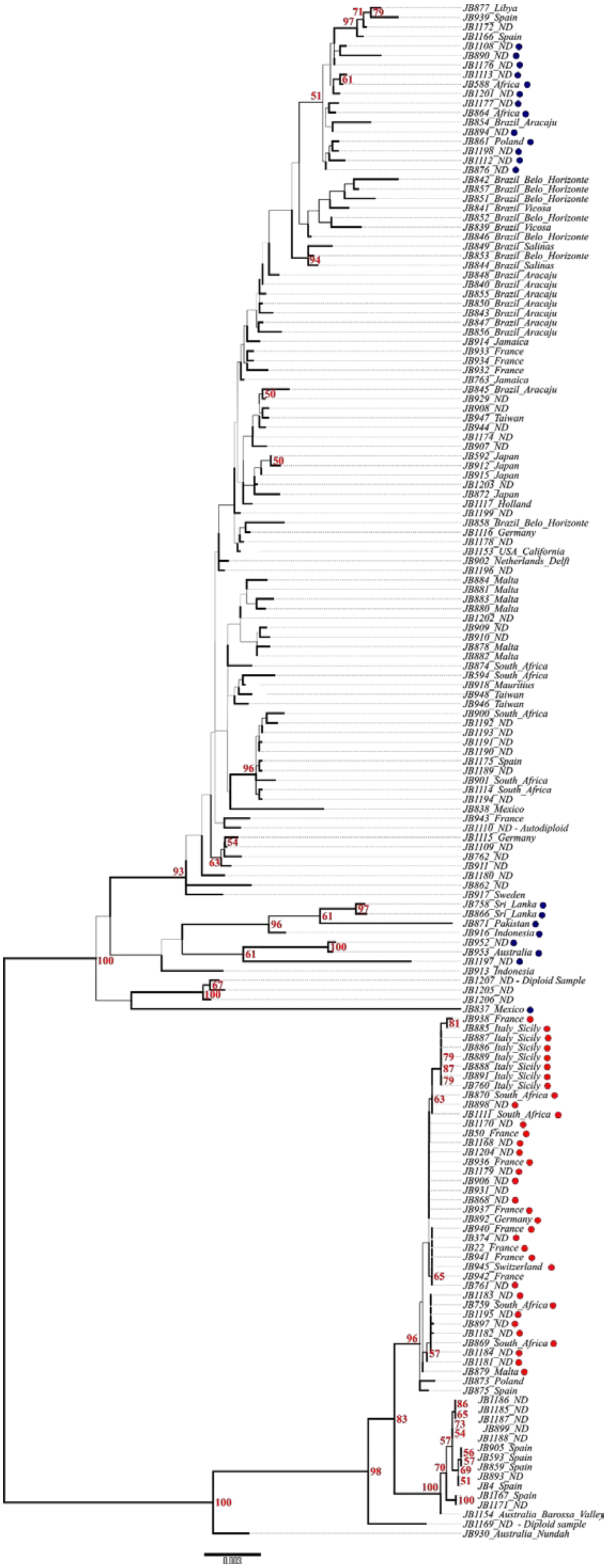
JB877_Libya
JB939_Spain
JB1172_ND
JB1166_Spain
JB1108_ND
JB890_ND
JB1176_ND
JB1113_ND
JB588_Africa
JB1201_ND
JB1177_ND
JB864_Africa
JB854_Brazil_Aracaju
JB894_ND
JB861_Poland
JB1198_ND
JB1112_ND
JB876_ND
JB842_Brazil_Belo_Horizonte
JB857_Brazil_Belo_Horizonte
JB851_Brazil_Belo_Horizonte
JB841_Brazil_Vicosa
JB852_Brazil_Belo_Horizonte
JB839_Brazil_Vicosa
JB846_Brazil_Belo_Horizonte
JB849_Brazil_Salinas
JB853_Brazil_Belo_Horizonte
JB844_Brazil_Salinas
JB848_Brazil_Aracaju
JB840_Brazil_Aracaju
JB855_Brazil_Aracaju
JB850_Brazil_Aracaju
JB843_Brazil_Aracaju
JB847_Brazil_Aracaju
JB856_Brazil_Aracaju
JB914_Jamaica
JB933_France
JB934_France
JB932_France
JB763_Jamaica
JB845_Brazil_Aracaju
JB929_ND
JB908_ND
JB947_Taiwan
JB944_ND
JB1174_ND
JB907_ND
JB592_Japan
JB912_Japan
JB915_Japan
JB1203_ND
JB872_Japan
JB1117_Holland
JB1199_ND
JB858_Brazil_Belo_Horizonte
JB1116_Germany
JB1178_ND
JB1153_USA_California
JB902_Netherlands_Delft
JB1196_ND
JB884_Malta
JB881_Malta
JB883_Malta
JB880_Malta
JB1202_ND
JB909_ND
JB910_ND
JB878_Malta
JB882_Malta
JB874_South_Africa
JB594_South_Africa
JB918_Mauritius
JB948_Taiwan
JB946_Taiwan
JB900_South_Africa
JB1192_ND
JB1193_ND
JB1191_ND
JB1190_ND
JB1175_Spain
JB1189_ND
JB901_South_Africa
JB1114_South_Africa
JB1194_ND
JB838_Mexico
JB943_France
JB1110_ND - Autodiploid
JB1115_Germany
JB1109_ND
JB762_ND
JB911_ND
JB1180_ND
JB862_ND
JB917_Sweden
JB758_Sri_Lanka
JB866_Sri_Lanka
JB871_Pakistan
JB916_Indonesia
JB952_ND
JB953_Australia
JB1197_ND
JB913_Indonesia
JB1207_ND - Diploid Sample
JB1205_ND
JB1206_ND
JB837_Mexico
JB938_France
JB885_Italy_Sicily
JB887_Italy_Sicily
JB886_Italy_Sicily
JB889_Italy_Sicily
JB888_Italy_Sicily
JB891_Italy_Sicily
JB760_Italy_Sicily
JB870_South_Africa
JB898_ND
JB1111_South_Africa
JB1170_ND
JB50_France
JB1168_ND
JB1204_ND
JB936_France
JB1179_ND
JB906_ND
JB931_ND
JB868_ND
JB937_France
JB892_Germany
JB940_France
JB374_ND
JB22_France
JB941_France
JB945_Switzerland
JB942_France
JB761_ND
JB1183_ND
JB759_South_Africa
JB1195_ND
JB897_ND
JB1182_ND
JB869_South_Africa
JB1184_ND
JB1181_ND
JB879_Malta
JB873_Poland
JB875_Spain
JB1186_ND
JB1185_ND
JB1187_ND
JB899_ND
JB1188_ND
JB905_Spain
JB593_Spain
JB859_Spain
JB893_ND
JB4_Spain
JB1167_Spain
JB1171_ND
JB1154_Australia_Barossa_Valley
JB1169_ND - Diploid sample
JB930_Australia_Nundah

**Supplementary Figure 8: Mitochondrial phylogeny.** Unrooted maximum-likelihood tree for 161 samples based on mitochondrial SNPs. Red numbers show bootstrapping support when higher than 50. Tips show sample ID and sampling location as in Supplementary Table 1. Scale bar units in substitutions per site. Samples with high proportion from one ancestral population are indicated with dots (red: high *Sp* clade proportion, blue: high *Sk* clade proportion).

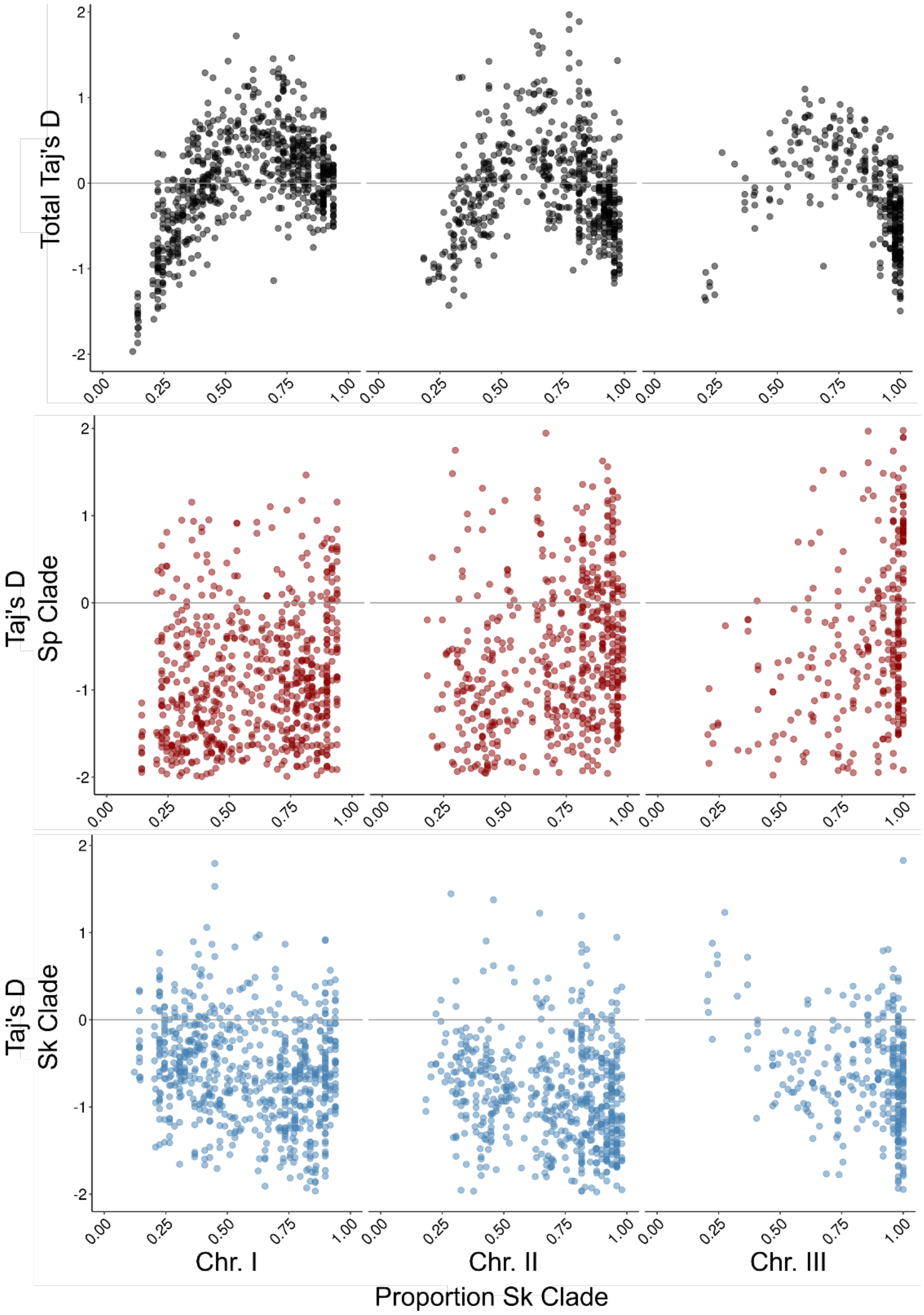


**Supplementary Figure 9: Tajima's D estimate.** Relationship between proportion of ancestral haplotypes (here represented by the proportion of *Sk* clade) and Tajima's D. Different panes show total Tajima's D (using all samples – black dots), and restricted to the *Sp* (red) or *Sk* (blue) clade.

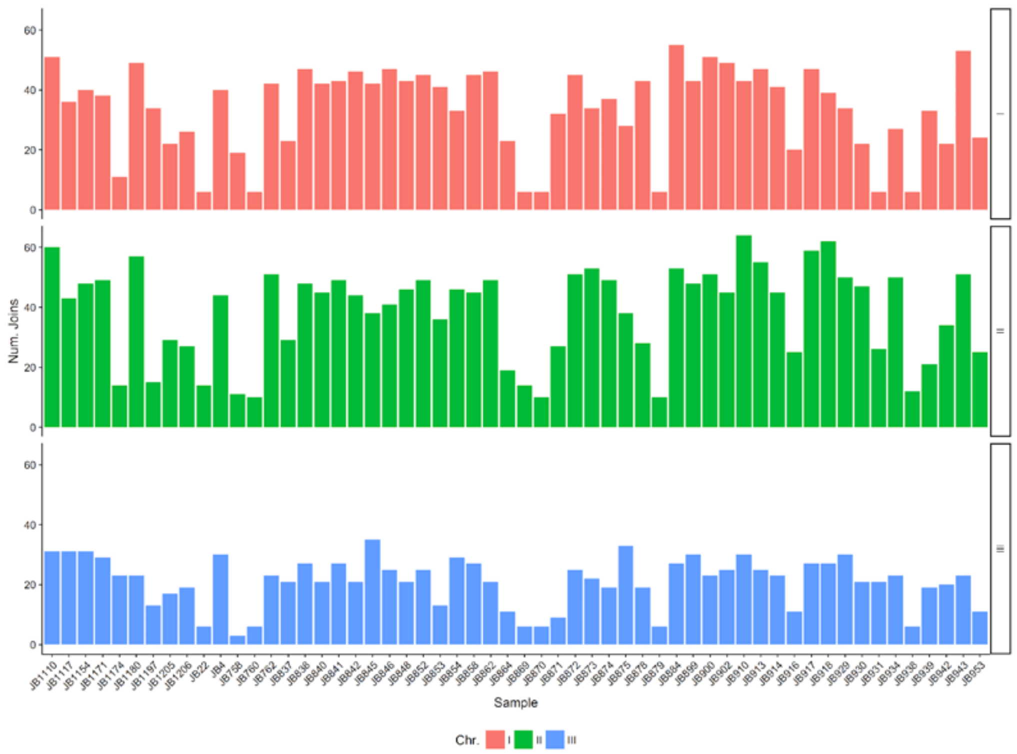


**Supplementary Figure 10: Number of join points per sample.** Number of transitions between *Sp* and *Sk* haplotypes along the genome per chromosome and sample.

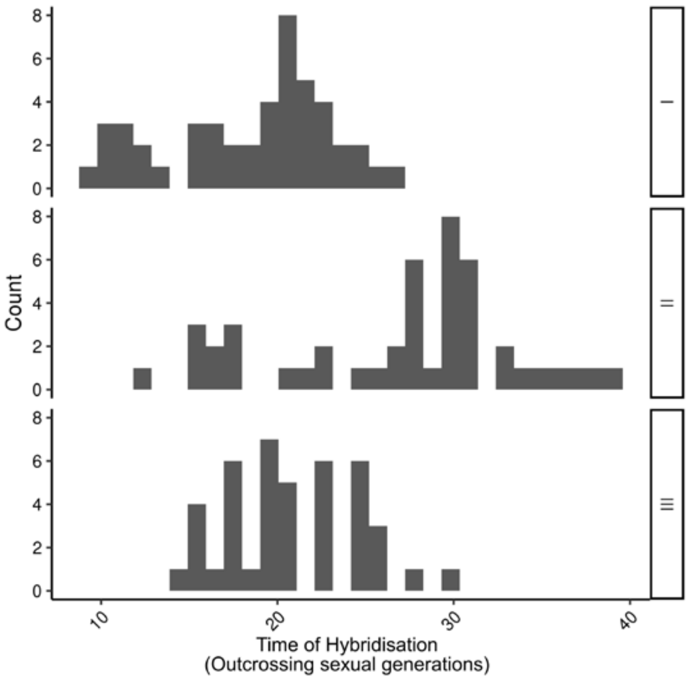


**Supplementary Figure 11: Estimated number of sexual generation since hybridisation.** Histogram of number of outcrossing sexual generation since hybridisation using the model by Janzen *et al.* (2018). Analysis was divided by chromosome and sample. Counts represent number of samples.

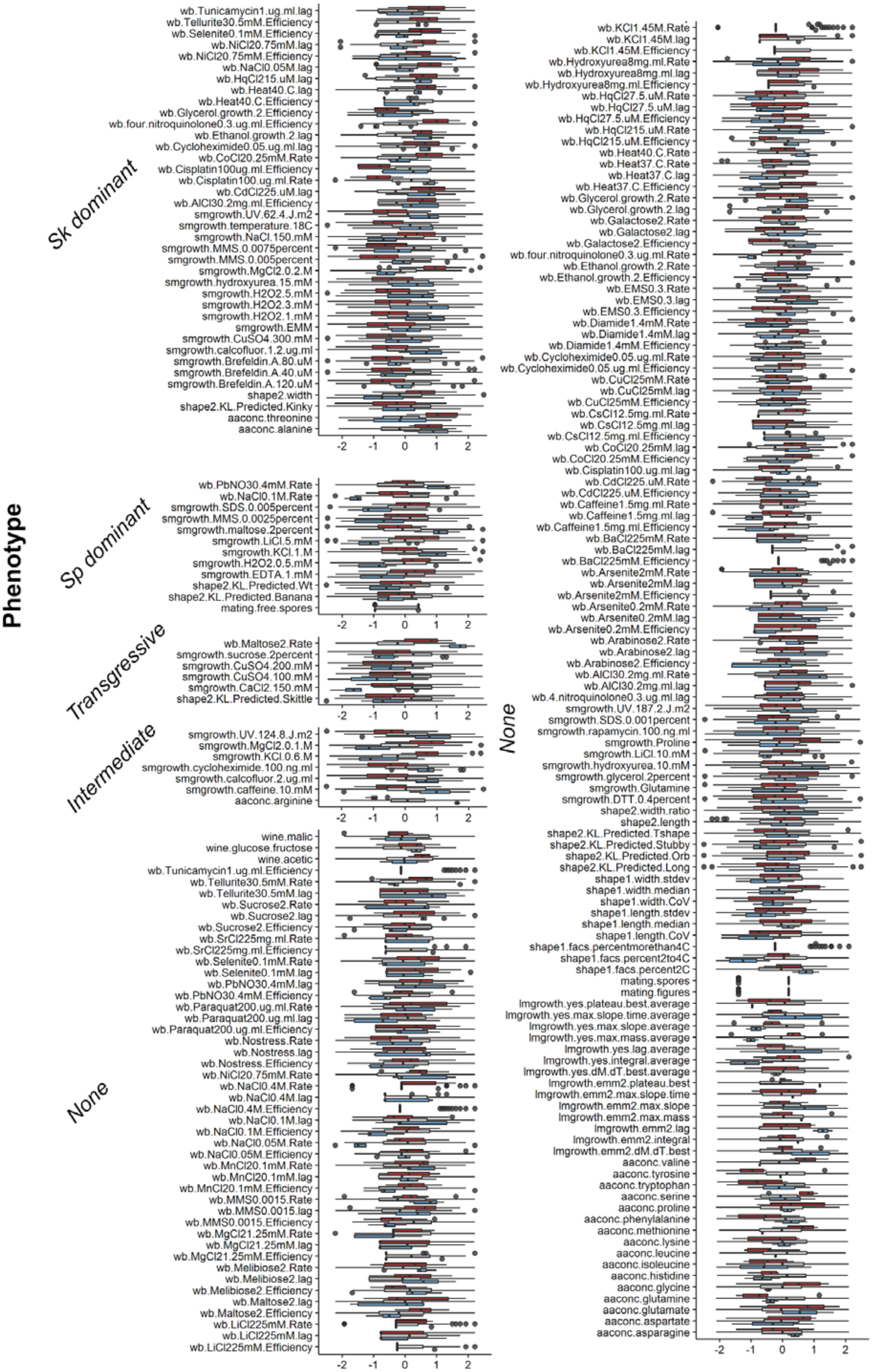

Phenotype
Sk dominant
wb.Tunicamycin1.ug.ml.lag
wb.Tellurite30.5mM.Efficiency
wb.Selenite0.1mM.Efficiency
wb.NiCl20.75mM.lag
wb.NiCl20.75mM.Efficiency
wb.NaCl0.05M.lag
wb.HqCl215.uM.lag
wb.Heat40.C.lag
wb.Heat40.C.Efficiency
wb.Glycerol.growth.2.Efficiency
wb.four.nitroquinolone0.3.ug.ml.Efficiency
wb.Ethanol.growth.2.lag
wb.Cycloheximide0.05.ug.ml.lag
wb.CoCl20.25mM.Rate
wb.Cisplatin100ug.ml.Efficiency
wb.Cisplatin100.ug.ml.Rate
wb.CdCl225.uM.lag
wb.AlCl30.2mg.ml.Efficiency
smgrowth.UV.62.4.J.m2
smgrowth.temperature.18C
smgrowth.NaCl.150.mM
smgrowth.MMS.0.0075percent
smgrowth.MMS.0.005percent
smgrowth.MgCl2.0.2.M
smgrowth.hydroxyurea.15.mM
smgrowth.H2O2.5.mM
smgrowth.H2O2.3.mM
smgrowth.H2O2.1.mM
smgrowth.EMM
smgrowth.CuSO4.300.mM
smgrowth.calcofluor.1.2.ug.ml
smgrowth.Brefeldin.A.80.uM
smgrowth.Brefeldin.A.40.uM
smgrowth.Brefeldin.A.120.uM
shape2.width
shape2.KL.Predicted.Kinky
aaconc.threonine
aaconc.alanine
-2 -1 0 1 2
Sp dominant
wb.PbNO30.4mM.Rate
wb.NaCl0.1M.Rate
smgrowth.SDS.0.005percent
smgrowth.MMS.0.0025percent
smgrowth.maltose.2percent
smgrowth.LiCl.5.mM
smgrowth.KCl.1.M
smgrowth.H2O2.0.5.mM
smgrowth.EDTA.1.mM
shape2.KL.Predicted.Wt
shape2.KL.Predicted.Banana
mating.free.spores
-2 -1 0 1 2
Transgressive
wb.Maltose2.Rate
smgrowth.sucrose.2percent
smgrowth.CuSO4.200.mM
smgrowth.CuSO4.100.mM
smgrowth.CaCl2.150.mM
shape2.KL.Predicted.Skittle
-2 -1 0 1 2
Intermediate
smgrowth.UV.124.8.J.m2
smgrowth.MgCl2.0.1.M
smgrowth.KCl.0.6.M
smgrowth.cycloheximide.100.ng.ml
smgrowth.calcofluor.2.ug.ml
smgrowth.caffeine.10.mM
aaconc.arginine
-2 -1 0 1 2
None
wine.malic
wine.glucose.fructose
wine.acetic
wb.Tunicamycin1.ug.ml.Efficiency
wb.Tellurite30.5mM.Rate
wb.Tellurite30.5mM.lag
wb.Sucrose2.Rate
wb.Sucrose2.lag
wb.Sucrose2.Efficiency
wb.SrCl225mg.ml.Rate
wb.SrCl225mg.ml.Efficiency
wb.Selenite0.1mM.Rate
wb.Selenite0.1mM.lag
wb.PbNO30.4mM.lag
wb.PbNO30.4mM.Efficiency
wb.Paraquat200.ug.ml.Rate
wb.Paraquat200.ug.ml.lag
wb.Paraquat200.ug.ml.Efficiency
wb.Nostress.Rate
wb.Nostress.lag
wb.Nostress.Efficiency
wb.NiCl20.75mM.Rate
wb.NaCl0.4M.Rate
wb.NaCl0.4M.lag
wb.NaCl0.4M.Efficiency
wb.NaCl0.1M.lag
wb.NaCl0.1M.Efficiency
wb.NaCl0.05M.Rate
wb.NaCl0.05M.Efficiency
wb.MnCl20.1mM.Rate
wb.MnCl20.1mM.lag
wb.MnCl20.1mM.Efficiency
wb.MMS0.0015.Rate
wb.MMS0.0015.lag
wb.MMS0.0015.Efficiency
wb.MgCl21.25mM.Rate
wb.MgCl21.25mM.lag
wb.MgCl21.25mM.Efficiency
wb.Melibiose2.Rate
wb.Melibiose2.lag
wb.Melibiose2.Efficiency
wb.Maltose2.lag
wb.Maltose2.Efficiency
wb.LiCl225mM.Rate
wb.LiCl225mM.lag
wb.LiCl225mM.Efficiency
-2 -1 0 1 2
None
wb.KCl1.45M.Rate
wb.KCl1.45M.lag
wb.KCl1.45M.Efficiency
wb.Hydroxyurea8mg.ml.Rate
wb.Hydroxyurea8mg.ml.lag
wb.Hydroxyurea8mg.ml.Efficiency
wb.HqCl27.5.uM.Rate
wb.HqCl27.5.uM.lag
wb.HqCl27.5.uM.Efficiency
wb.HqCl215.uM.Rate
wb.HqCl215.uM.Efficiency
wb.Heat40.C.Rate
wb.Heat37.C.Rate
wb.Heat37.C.lag
wb.Heat37.C.Efficiency
wb.Glycerol.growth.2.Rate
wb.Glycerol.growth.2.lag
wb.Galactose2.Rate
wb.Galactose2.lag
wb.Galactose2.Efficiency
wb.four.nitroquinolone0.3.ug.ml.Rate
wb.Ethanol.growth.2.Rate
wb.Ethanol.growth.2.Efficiency
wb.EMS0.3.Rate
wb.EMS0.3.lag
wb.EMS0.3.Efficiency
wb.Diamide1.4mM.Rate
wb.Diamide1.4mM.lag
wb.Diamide1.4mM.Efficiency
wb.Cycloheximide0.05.ug.ml.Rate
wb.Cycloheximide0.05.ug.ml.Efficiency
wb.CuCl25mM.Rate
wb.CuCl25mM.lag
wb.CuCl25mM.Efficiency
wb.CsCl12.5mg.ml.Rate
wb.CsCl12.5mg.ml.lag
wb.CsCl12.5mg.ml.Efficiency
wb.CoCl20.25mM.lag
wb.CoCl20.25mM.Efficiency
wb.Cisplatin100.ug.ml.lag
wb.CdCl225.uM.Rate
wb.CdCl225.uM.Efficiency
wb.Caffeine1.5mg.ml.Rate
wb.Caffeine1.5mg.ml.lag
wb.Caffeine1.5mg.ml.Efficiency
wb.BaCl225mM.Rate
wb.BaCl225mM.lag
wb.BaCl225mM.Efficiency
wb.Arsenite2mM.Rate
wb.Arsenite2mM.lag
wb.Arsenite2mM.Efficiency
wb.Arsenite0.2mM.Rate
wb.Arsenite0.2mM.lag
wb.Arsenite0.2mM.Efficiency
wb.Arabinose2.Rate
wb.Arabinose2.lag
wb.Arabinose2.Efficiency
wb.AlCl30.2mg.ml.Rate
wb.AlCl30.2mg.ml.lag
wb.4.nitroquinolone0.3.ug.ml.lag
smgrowth.UV.187.2.J.m2
smgrowth.SDS.0.001percent
smgrowth.rapamycin.100.ng.ml
smgrowth.Proline
smgrowth.LiCl.10.mM
smgrowth.hydroxyurea.10.mM
smgrowth.glycerol.2percent
smgrowth.Glutamine
smgrowth.DTT.0.4percent
shape2.width.ratio
shape2.length
shape2.KL.Predicted.Tshape
shape2.KL.Predicted.Stubby
shape2.KL.Predicted.Orb
shape2.KL.Predicted.Long
shape1.width.stdev
shape1.width.median
shape1.width.CoV
shape1.length.stdev
shape1.length.median
shape1.length.CoV
shape1.facs.percentmorethan4C
shape1.facs.percent2to4C
shape1.facs.percent2C
mating.spores
mating.figures
lmgrowth.yes.plateau.best.average
lmgrowth.yes.max.slope.time.average
lmgrowth.yes.max.slope.average
lmgrowth.yes.max.mass.average
lmgrowth.yes.lag.average
lmgrowth.yes.integral.average
lmgrowth.yes.dM.dT.best.average
lmgrowth.emm2.plateau.best
lmgrowth.emm2.max.slope.time
lmgrowth.emm2.max.slope
lmgrowth.emm2.max.mass
lmgrowth.emm2.lag
lmgrowth.emm2.integral
lmgrowth.emm2.dM.dT.best
aaconc.valine
aaconc.tyrosine
aaconc.tryptophan
aaconc.serine
aaconc.proline
aaconc.phenylalanine
aaconc.methionine
aaconc.lysine
aaconc.leucine
aaconc.isoleucine
aaconc.histidine
aaconc.glycine
aaconc.glutamine
aaconc.glutamate
aaconc.aspartate
aaconc.asparagine
-2 -1 0 1 2

**Supplementary Figure 12: Phenotypic distribution per trait.** Boxplot of phenotypic distribution using normalised values. Traits are divided by ancestral admixture proportions: pure *Sk* clade (blue), pure *Sp* clade (red) and hybrids (gray).

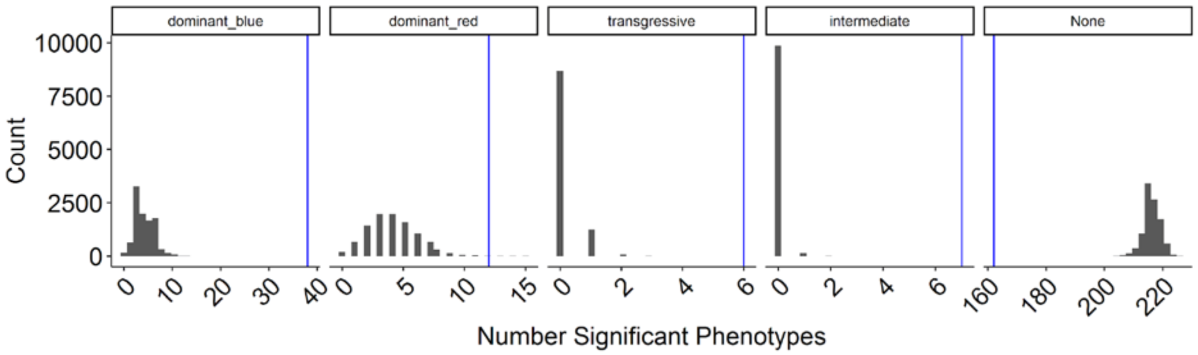


**Supplementary Figure 13: Permutation test with phenotypic data.** Histogram of total number of significant traits after randomising sample category (pure *Sp* clade , pure *Sk* clade and hybrids) without replacement. Blue lines show observed number of significant traits. Matrix data was randomised 10000 times to produce distribution per category group.

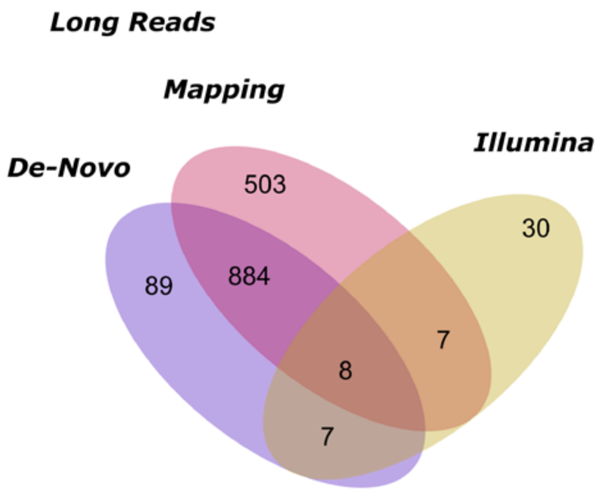


**Supplementary Figure 14: Total number of structural variant calls.** Venn diagram comparing total number of structural variants in reported list from Illumina reads (Jeffares, 2017), and long reads. The last one is divided between the two genotype calling approaches: *de-novo* and mapping reads.

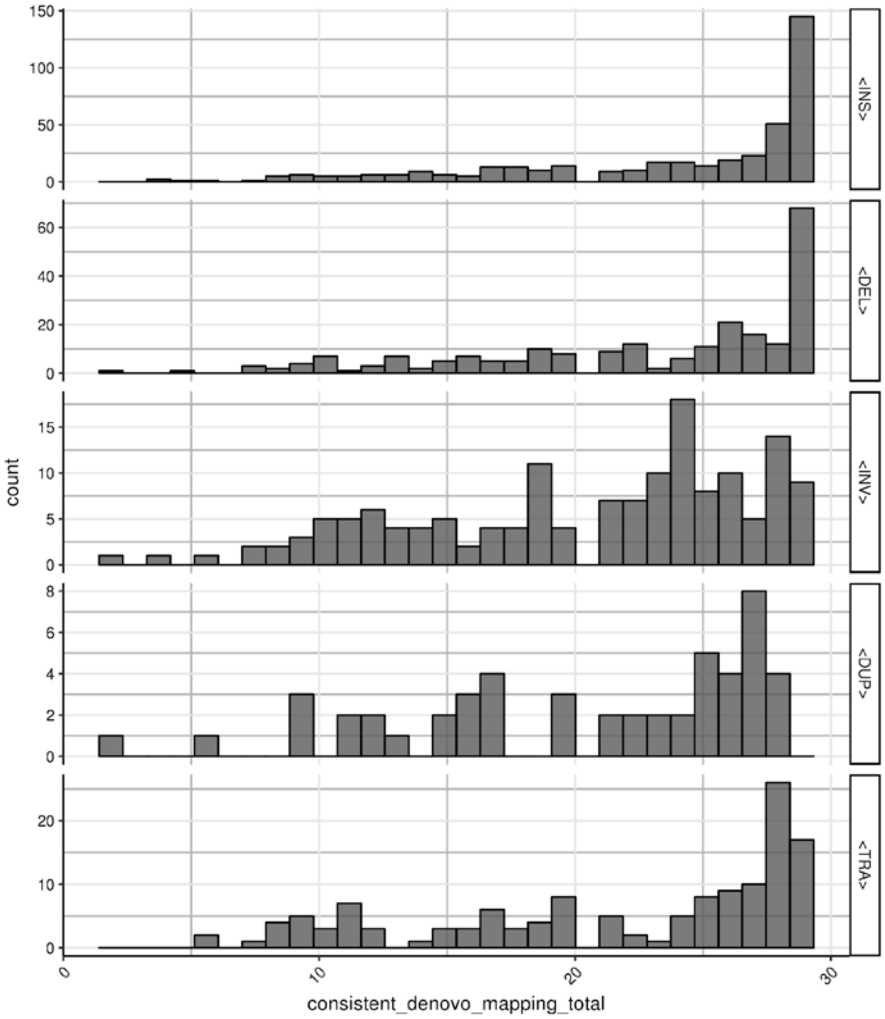


**Supplementary Figure 15: Comparison between SV detection using the de-novo vs. mapping approach.** Histogram of consistent genotypes of SVs in samples sequenced with PacBio and/or Nanopore (29 in total). Planes are divided by type of SV (INS: insertion, DEL: deletion, INV: inversion, DUP: duplication, TRA: translocation). The obtained genotype from the de-novo and mapping approach in each sample and variant was compared. The histogram shows the number of variants with consistent mapping and de-novo genotype (no necessarily the same genotype between samples). For example, there are around 150 insertions with consistent mapping and de-novo genotype in all 29 samples. In another 50 insertion one sample out of the total shows different genotype between the mapping and de-novo approach.

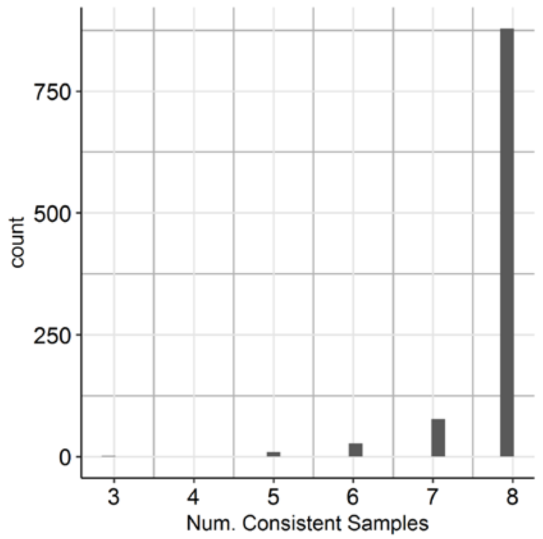


**Supplementary Figure 16: Consistency of SV genotype detection using PacBio vs. MinIon data.** Histogram of consistent genotypes of SVs in samples sequenced with both PacBio and Nanopore (8 strains in total). For each variants the genotype between the PacBio and Nanopore data was compared in each sample. The histogram shows the number of variants with up to 8 consistent samples. For example, 878 SVs show the same genotype per sample between PacBio and Nanopore data (no necessarily the same genotype between samples).

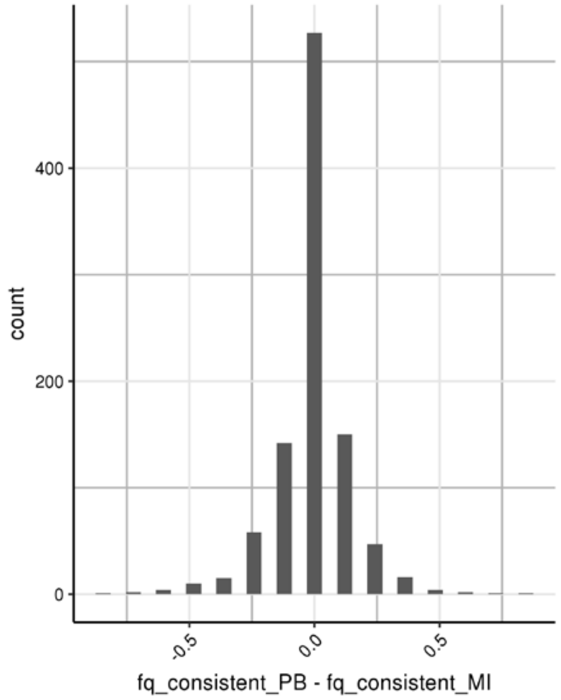


**Supplementary Figure 17: Difference allele frequency using PacBio vs. MinIon data.** Histogram of difference in observed allele frequency of samples sequenced with both PacBio and Nanopore (8 in total).

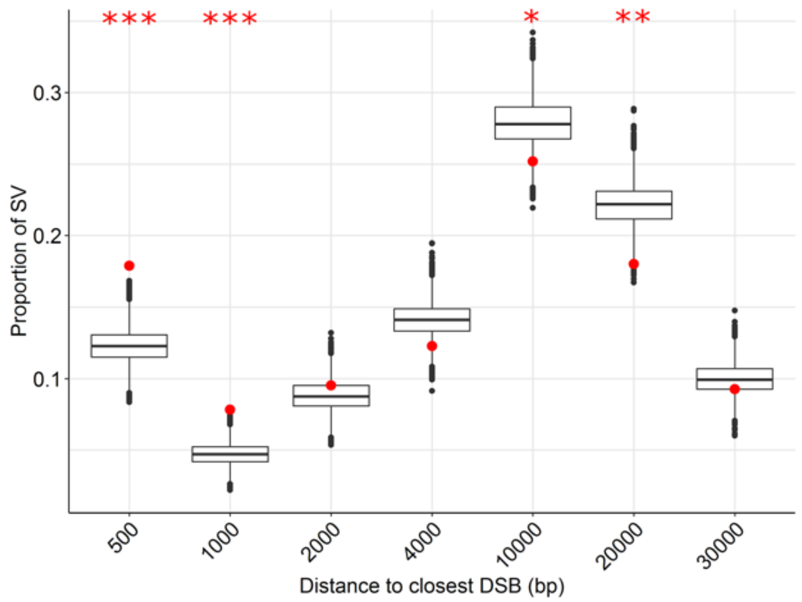


**Supplementary Figure 18: Boxplot displaying the proportion of SV relative to the physical genetic distance from the closest double-strand break.** Boxplot with the distribution of the expected proportion of structural variants by random permutation within different genetic distances. SV observed within 500bp and 1kb of the closes DSB are significantly overrepresented and those at 10 or 20kb are underrepresented. Red points represent the observed proportions from long read sequencing data. Significance in difference between permutations and observed values are shown with asterisks: * p-value < 0.05, ** p-value <0.001, *** p-value <0.0001.

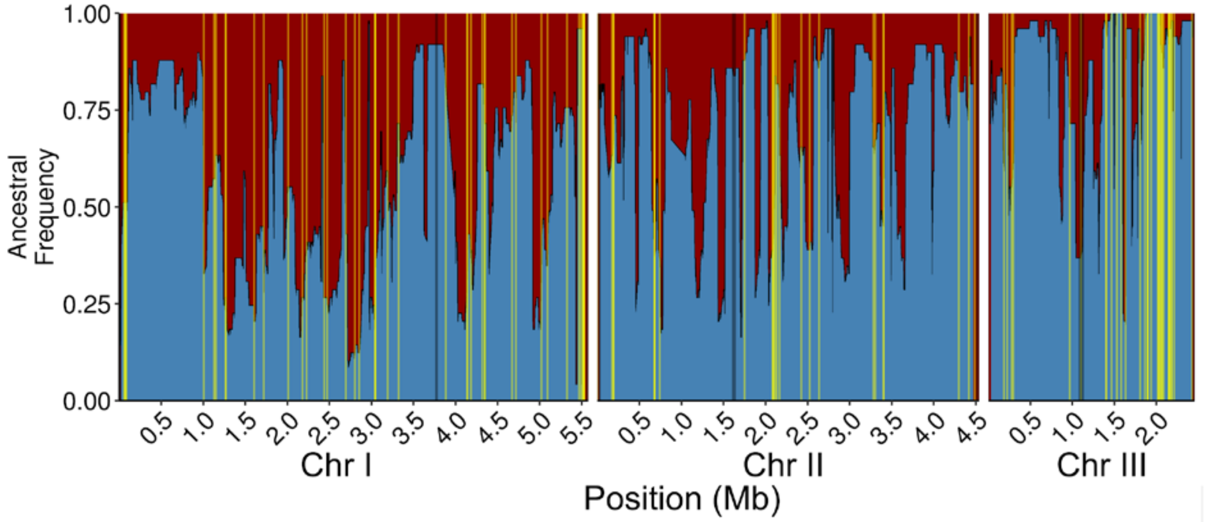


**Supplementary Figure 19: Distribution of divergent SV.** Proportion of *Sp* (red) and *Sk* (blue) ancestry across all 57 samples along the genome. Divergent between ancestral groups SV (with frequency difference > 0.7 between *Sp* and *Sk*) are shown with yellow lines. Centromeric regions are shown in gray.

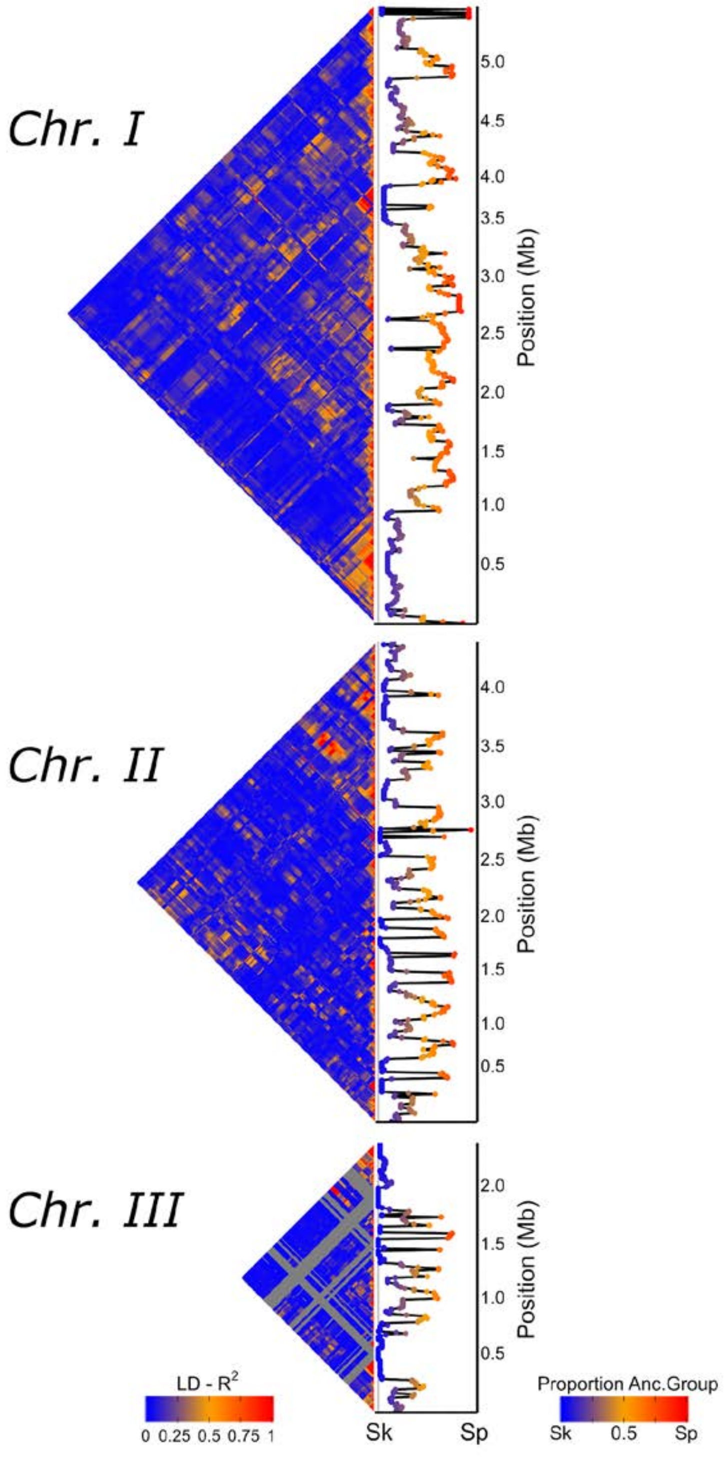


**Supplementary Figure 20: Linkage disequilibrium between pairs of genomic windows polarized by ancestry.** Heat map of linkage disequilibrium ($R^2$) for all comparison between genomic windows (left panel). Proportions of ancestral groups (*Sp* or *Sk*) along the genome are shown in the right panel. Genomic regions fixed for one of the ancestral groups are shown in gray areas in the heat map.

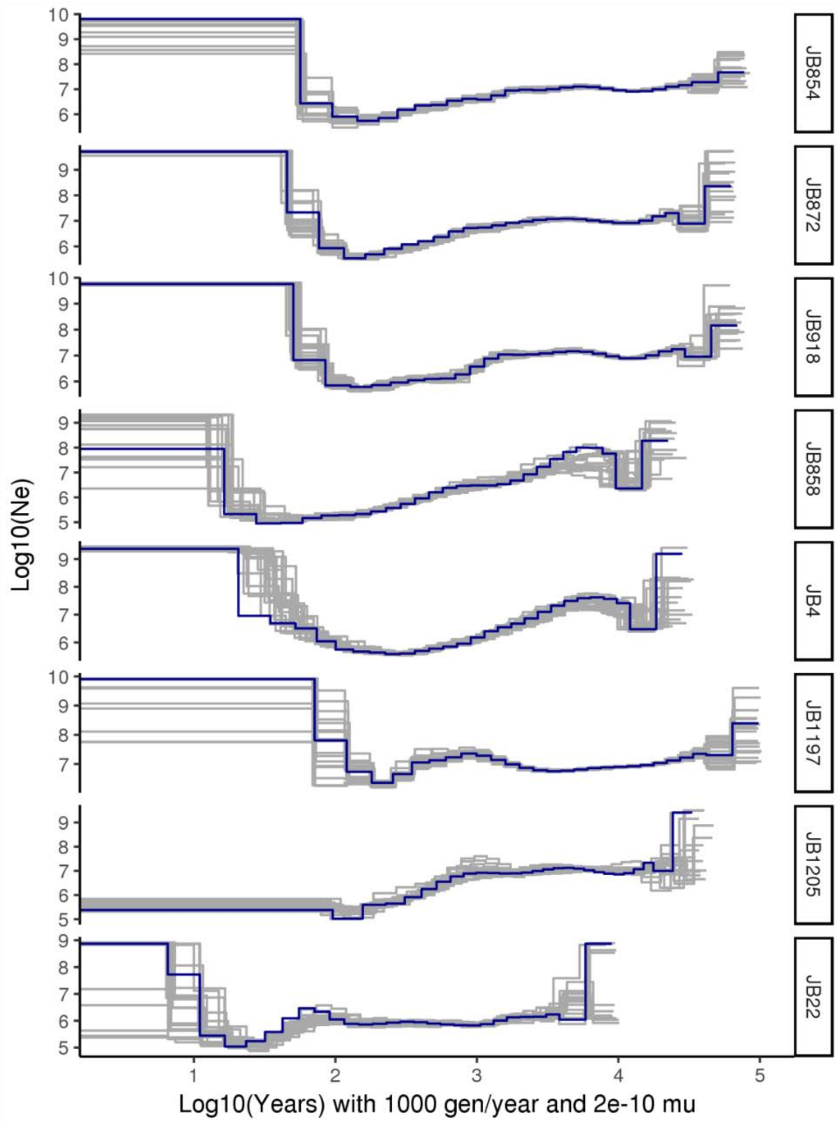


**Supplementary Figure 21: Inferred effective population size over time with MSMC.** Analysis was divided by secondary clusters presented in Figure 1c. Each cluster is label with one representative of the cluster. Inferred values are shown with the blue line and bootstraps with grey lines. Time in x axis in log10 scale using 1000 generations per year and mutation rate of $2x10^{-10}$. Present time is in x=0 and increase in the past. Effective population size in log10 scale.

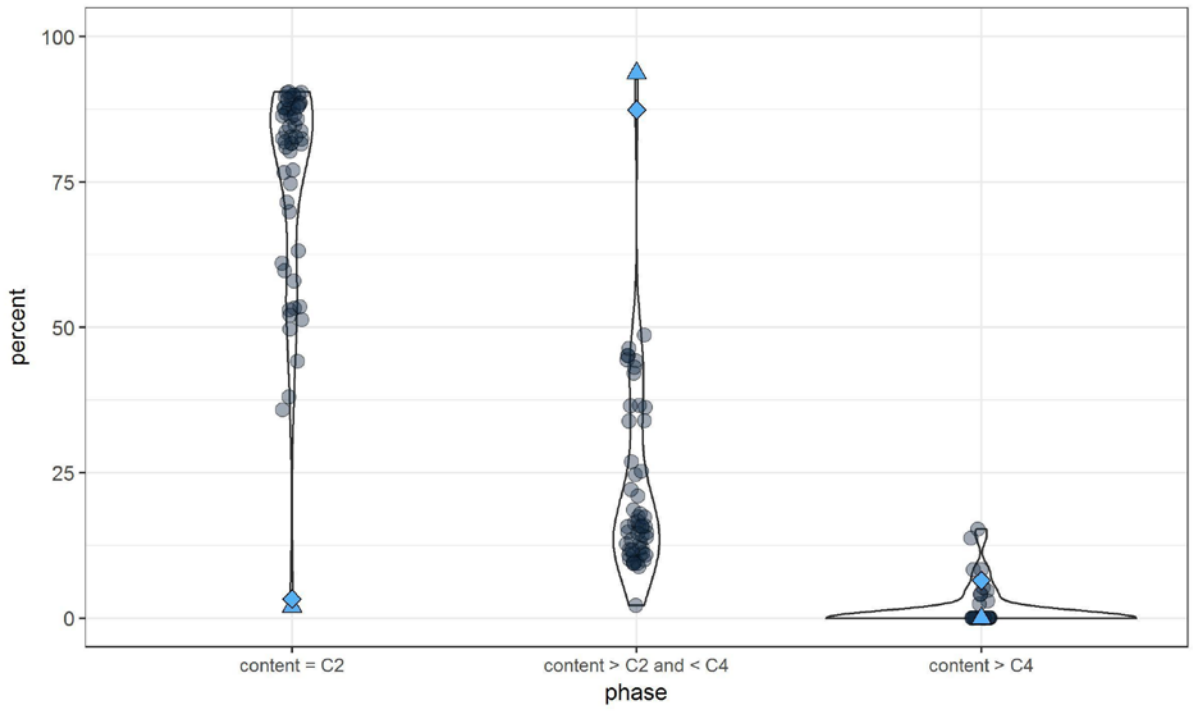


**Supplementary figure 22: DNA content per strain.** Representation of flow cytometry measurements of percentage of cells with nuclear DNA content <C2 (two times haploid genomic content; left), nuclear content between C2 and C4 (middle) and larger than C4 (right). Due to the short G1 phase and late cytokinesis, few cells with C1 content are found in fission yeast. Haploid cells are thus found in mostly in the C2 phase. Diploid cells will be mostly in C4 or larger. The strain with high heterozygosity (JB1207) is indicated by the blue diamond and it has a large proportion of cells with large genomic content. Additionally, strain JB1169 (blue triangle) appears to be of higher ploidy, though it does not show increased heterozygosity, which suggests this strain was formed by auto-diploidization. Data from Jeffares et al. 2015.

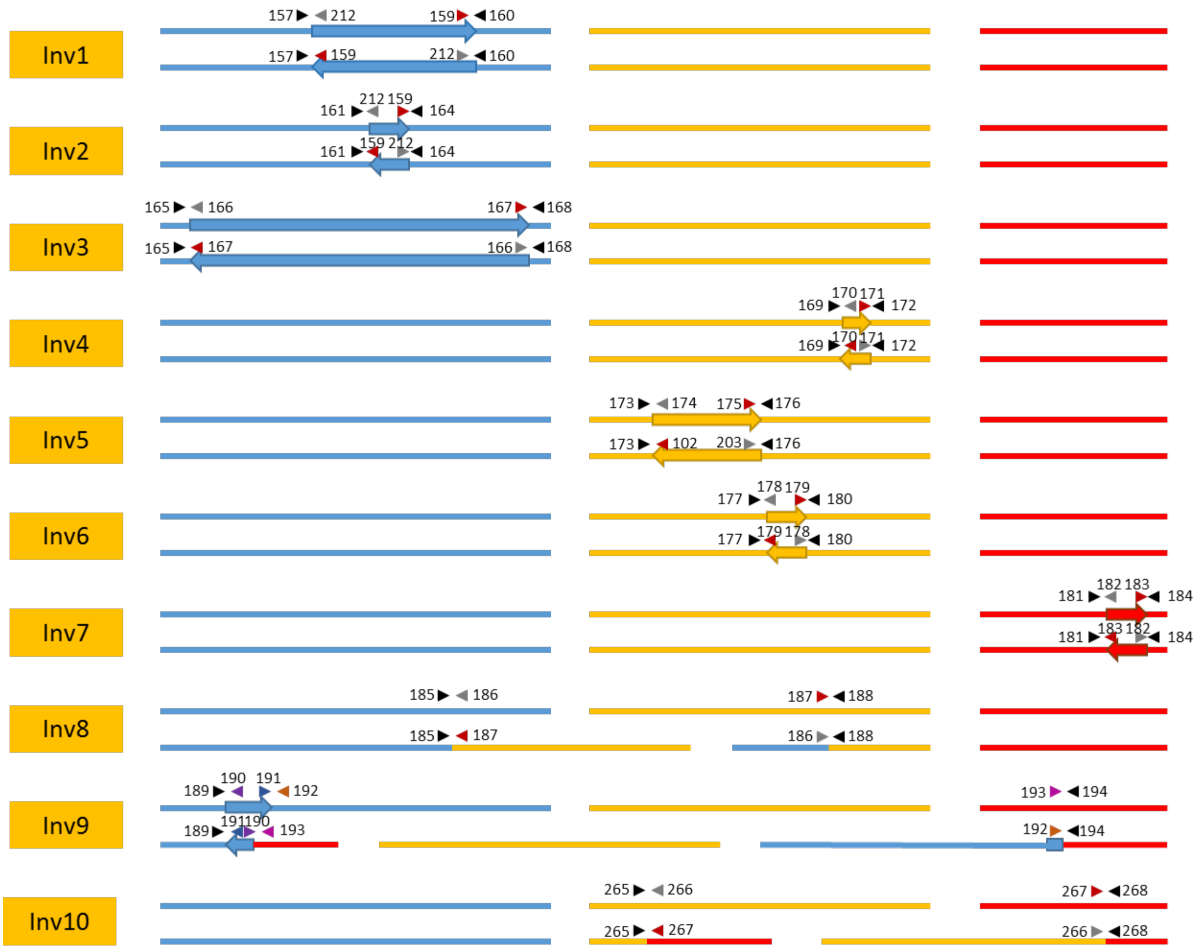


**Supplementary Figure 23: Overview of PCR primers used to verify inversions and translocations.** For each SV first the positions in the reference genome are given (top), and below the organization in the rearranged form as observed in the long-read *de novo* assemblies. The three lines represent Chromosomes I, II and III. The small arrows indicate the position and direction of the primers with the numbers of the primers indicated. The large arrows indicate the size and orientation of inversions. Note that more than one SV can occur per strain, and that the representations are thus not always representations of actual strains. Primers in ‘Inv10’ from Zanders et al. 2014.
